# DprA recruits ComM to facilitate recombination during natural transformation in Gram-negative bacteria

**DOI:** 10.1101/2024.10.21.619469

**Authors:** Triana N. Dalia, Mérick Machouri, Céline Lacrouts, Yoann Fauconnet, Raphaël Guerois, Jessica Andreani, J. Pablo Radicella, Ankur B. Dalia

## Abstract

Natural transformation (NT) represents one of the major modes of horizontal gene transfer in bacterial species. During NT, cells can take up free DNA from the environment and integrate it into their genome by homologous recombination. While NT has been studied for >90 years, the molecular details underlying this recombination remain poorly understood. Recent work has demonstrated that ComM is an NT-specific hexameric helicase that promotes recombinational branch migration in Gram-negative bacteria. How ComM is loaded onto the post-synaptic recombination intermediate during NT, however, remains unclear. Another NT-specific recombination mediator protein that is ubiquitously conserved in both Gram-positive and Gram-negative bacteria is DprA. Here, we uncover that DprA homologs in Gram-negative species contain a C-terminal winged helix domain that is predicted to interact with ComM by AlphaFold. Using *Helicobacter pylori* and *Vibrio cholerae* as model systems, we demonstrate that ComM directly interacts with the DprA winged-helix domain, and that this interaction is critical for DprA to recruit ComM to the recombination site to promote branch migration during NT. These results advance our molecular understanding of recombination during this conserved mode of horizontal gene transfer. Furthermore, they demonstrate how structural modeling can help uncover unexpected interactions between well-studied proteins to provide deep mechanistic insight into the molecular coordination required for their activity.

**SIGNIFICANCE STATEMENT:** Bacteria can acquire novel traits like antibiotic resistance and virulence through horizontal gene transfer by natural transformation. During this process, cells take up free DNA from the environment and integrate it into their genome by homologous recombination. Many of the molecular details underlying this process, however, remain incompletely understood. In this study, we identify a new protein-protein interaction between ComM and DprA, two factors that promote homologous recombination during natural transformation in Gram-negative species. Through a combination of bioinformatics, structural modeling, cell biological assays, and complementary genetic approaches, we demonstrate that this interaction is required for DprA to recruit ComM to the site of homologous recombination.

## INTRODUCTION

Natural transformation (NT) is a broadly conserved mechanism of horizontal gene transfer present in diverse Gram-positive and Gram-negative eubacteria, as well as some archaeal species^1,2^. During this process, cells translocate single-stranded transforming DNA (tDNA) across the cytoplasmic membrane^3–5^, which can then be integrated into the genome by homologous recombination. This can promote the acquisition of novel traits including antibiotic resistance and virulence factors in bacterial pathogens.

During NT, the primary bacterial recombinase, RecA, binds to tDNA and both (1) facilitates a homology search and (2) initiates the recombination reaction by promoting strand invasion of the bacterial chromosome to generate a 3-stranded D-loop structure. This 3-stranded intermediate is distinct from the more canonical recombination intermediates that form between two dsDNA substrates during DNA repair. For example, the 3-stranded intermediate during NT can never form a classical Holliday junction. As a result, the recombination reaction that occurs during NT must be distinct from more canonical forms of dsDNA recombination. Consistent with this, there are a number of proteins that have been characterized that are uniquely required for recombination during NT.

One of those proteins is DprA, which is present in all naturally transformable species^2^. DprA is a ssDNA-binding recombination mediator protein. During NT, DprA has been implicated in both (1) protecting ssDNA that has been internalized into the cytoplasm and (2) facilitating recombination by aiding in the recruitment of RecA^6–10^. Thus, to date, DprA has primarily been associated with the early pre-synaptic steps of recombination that lead to formation of the D-loop.

Another NT-specific recombination protein is ComM, which is only conserved in Gram-negative naturally transformable species^11^. ComM is a NT-specific helicase that binds to the crossover junctions (*i.e.*, branches) of the D-loop to expand them via recombinational branch migration^11,12^. This activity facilitates the integration of heterologous sequences during NT.

While most of the proteins involved in recombination during NT have been identified and characterized to some degree, the molecular coordination between these factors remains poorly understood. Here, we uncover that an extended C-terminal winged-helix domain of Gram-negative DprA homologs^13^ helps to recruit ComM to the site of tDNA recombination.

## RESULTS

### ComM-DprA interactions promote NT in Helicobacter pylori

DprA is broadly conserved in bacterial species and is always found in species capable of NT^2^. Prior work characterized the structure of the C-terminal winged helix domain of *Helicobacter pylori* DprA, which is present in a subset of DprA proteins^13^. The DprA_Hp_ C-terminal domain is not involved in DNA-binding despite displaying a winged helix fold^13,14^. Because this domain is required for efficient NT, it was proposed that its function could be to establish interactions with other NT-specific proteins^13^. We therefore analyzed DprA in a diverse set of bacterial species and uncovered that the DprA homologs among Gram-negative species commonly encode this extended C-terminal winged helix domain, but it is absent in the DprA homologs of Gram-positive species (**Fig. S1, Dataset S1**). This distribution largely mirrors the distribution of ComM, where species that contain a ComM homolog also encode a DprA homolog with a C-terminal winged-helix domain (**Fig. S1, Dataset S1**).

Thus, we hypothesized that the DprA winged-helix domain may directly interact with ComM to facilitate NT. To test this, we first used *H. pylori* as a model system. Consistent with our hypothesis, AlphaFold 3 (AF3)^15^ predicts a high-confidence interaction between the DprA_Hp_ winged-helix domain and ComM_Hp_ (ipTM = 0.76; pTM = 0.86)(**Fig. 1A**). To test whether these proteins interact empirically, we performed bacterial adenylate cyclase two-hybrid (BACTH) assays between the DprA_Hp_ ^C-ter^ (winged-helix domain) and full length ComM. In these assays, LacZ activity is directly linked to the interaction of these proteins^16^. This BACTH analysis revealed that DprA_Hp_ and ComM_Hp_ interact with one another (**Fig. 1B**, **Fig. S2**).

**Fig. 1.**
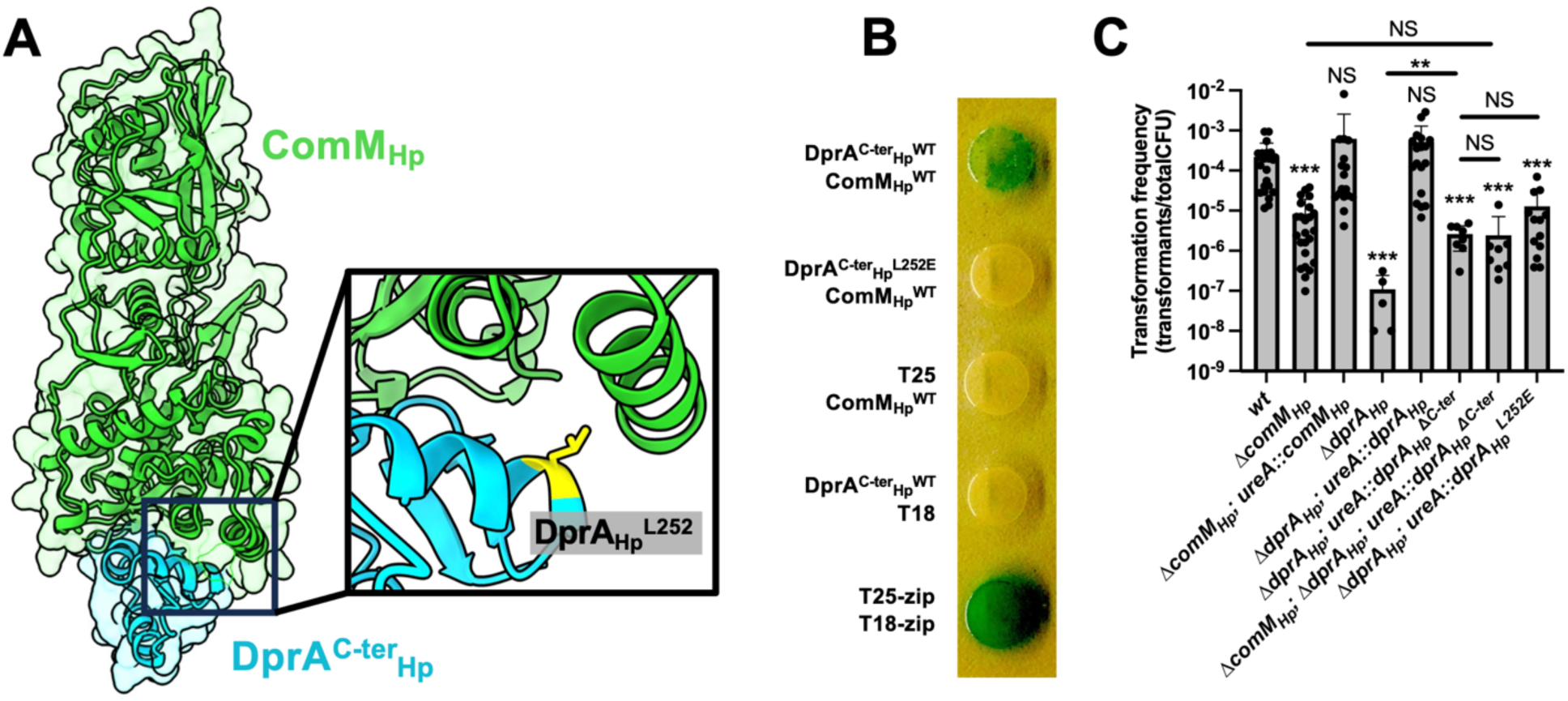
ComM-DprA interactions promote NT in *H. pylori*. (**A**) AlphaFold 3 model of ComM_Hp_-DprA^C-ter^_HP_ interactions. The inset shows the DprA_Hp_ ^L252^ interaction interface residue targeted for mutagenesis in yellow. (**B**) BACTH analysis to assess the interaction between the indicated alleles of DprA and ComM. All DprA constructs for BACTH consisted of C-terminal fusions of T25 to the terminal 53 amino acids of the indicated DprA_Hp_ allele. The ComM construct for BACTH consisted of a C-terminal fusion of T18 to full length ComM_Hp_. The T25-zip and T18-zip constructs represent a known interacting pair and are a positive control in this assay. Data are representative of two independent experiments. (**C**) Natural transformation assays of the indicated *H. pylori* strains using linear tDNA that integrates into the genome. Data are from at least 5 independent biological replicates and shown as the mean ± SD. Statistical comparisons were made by one-way ANOVA with Tukey’s multiple comparison test of the log-transformed data. NS, not significant. *** = *p* < 0.001. Statistical identifiers directly above bars represent comparisons to the *wt*.

To test whether the interaction between ComM_Hp_ and DprA_Hp_ is functionally relevant, we sought to assess the impact of disrupting these interactions on NT frequency in *H. pylori*. For these assays, *dprA_Hp_* and *comM_Hp_* alleles were expressed ectopically at the *ureA* locus. Importantly, ectopic expression of the wildtype alleles (*comM_Hp_* and *dprA_Hp_*) restored NT in the strains where the native gene was deleted (**Fig. 1C**). When cells were complemented with an allele of *dprA* lacking the C-terminal winged-helix domain (*dprA_Hp_ ^ΔC-ter^*) the NT frequency was indistinguishable from *ΔcomM_Hp_* (**Fig. 1C**). Importantly, *dprA_Hp_ ^ΔC-ter^* transformed at rates significantly higher than Δ*dprA*, suggesting that deleting the DprA_Hp_ winged-helix did not abolish all DprA activity. Thus, we hypothesized that the reduced NT frequency observed in *dprA_Hp_ ^ΔC-ter^* was due to ablation of DprA_Hp_-ComM_Hp_ interactions. To test this more rigorously, we assessed the phenotype of a *dprA_Hp_ ^ΔC-ter^ ΔcomM_Hp_* double mutant.

If the reduced NT observed in *dprA_Hp_ ^ΔC-ter^* is due to ablation of DprA_Hp_-ComM_Hp_ interactions, we hypothesized that these mutations should be epistatic. Indeed, we found that the phenotype of *dprA_Hp_ ^ΔC-ter^ ΔcomM_Hp_* double mutant was not significantly different from either the *dprA_Hp_ ^ΔC-ter^* or *ΔcomM_Hp_* single mutants (**Fig. 1C**). This result is consistent with DprA_Hp_-ComM_Hp_ interactions facilitating NT.

To test this further, we analyzed the AF3 model of DprA_Hp_-ComM_Hp_ interactions and found that DprA_Hp_ ^L252^ is buried within the interface of these two proteins (**Fig. 1A**). Thus, we hypothesized that introducing a charged amino acid at this site should disrupt DprA_Hp_-ComM_Hp_ interactions. Consistent with this, we found that DprA_Hp_ ^L252E^ no longer interacts with ComM_Hp_ in BACTH assays (**Fig. 1B**). Moreover, *dprA_Hp_ ^L252E^* phenocopies *dprA_Hp_ ^ΔC-ter^* in NT frequency assays (**Fig. 1C**). Interestingly, DprA_Hp_ ^L252A^ did not diminish interactions with ComM_Hp_ by BACTH (**Fig. S2**), which suggests that the charged glutamate at this position (DprA_Hp_ ^L252E^) is necessary to disrupt DprA_Hp_-ComM_Hp_ interactions. All together, these data strongly suggest that DprA_Hp_ - ComM_Hp_ interactions are critical for ComM_Hp_ activity in *H. pylori*.

### ComM-DprA interactions promote recombination of tDNA during NT in V. cholerae

The DprA C-terminal winged-helix domain is widely distributed among Gram-negative bacteria (**Fig. S1**). This includes *Vibrio cholerae*, the organism in which ComM was first characterized as a branch migration factor, and a species in which many tools have already been established for studying ComM activity^11,17^. So next, we sought to use *V. cholerae* as a model system to elucidate (1) whether ComM-DprA interactions are broadly conserved and if so, (2) how DprA-ComM interactions mechanistically promote NT.

First, we found that AF3 predicted a high confidence interaction between the DprA_Vc_ winged-helix domain and ComM_Vc_ (ipTM = 0.92; pTM = 0.85) (**Fig. 2A**). Consistent with this, we found that the DprA_Vc_ winged-helix domain and ComM_Vc_ interact with one another in BACTH assays (**Fig. 2B**).

**Fig. 2.**
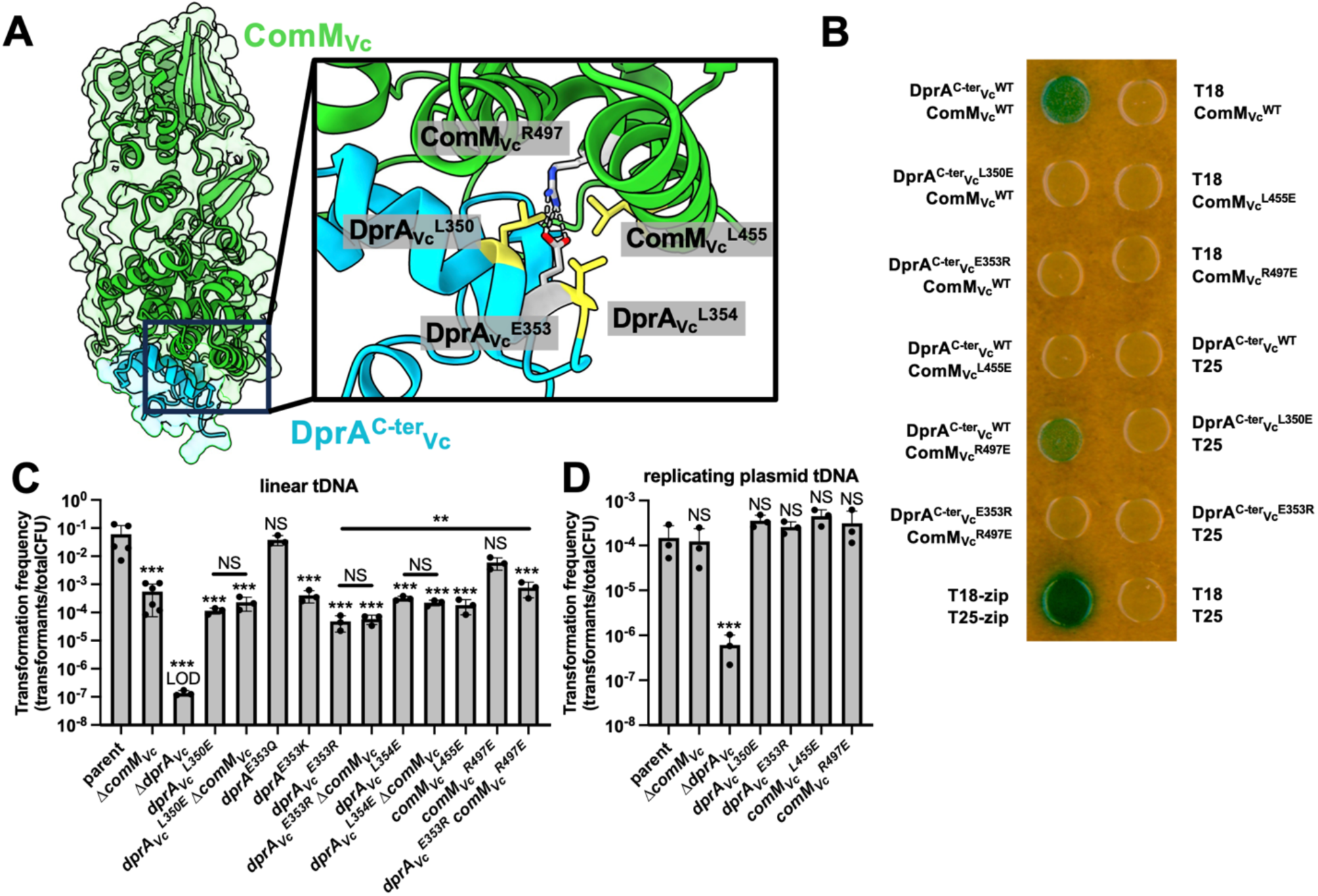
ComM-DprA interactions promote recombination of tDNA during NT in *V. cholerae*. (**A**) AlphaFold 3 model of ComM_Vc_-DprA^C-ter^_Vc_ interactions. The inset shows the interaction interface residues targeted for mutagenesis. The residues that form a putative electrostatic interaction (ComM_Vc_ ^R497^ and DprA_Vc_ ^E353^) are in gray with charged atoms in blue (positive) or red (negative), while the leucine residues within the interface (DprA_Vc_ ^L350^, DprA_Vc_ ^L354^, and ComM_Vc_ ^L455^) are colored yellow. (**B**) BACTH analysis to assess the interaction between the indicated alleles of DprA and ComM. All DprA constructs for BACTH consisted of N-terminal fusions of T18 to the terminal 71 amino acids of the indicated DprA allele. All ComM constructs for BACTH consisted of C-terminal fusions of T25 to full length ComM allele indicated. Data in **B** are representative of two independent experiments. (**C**) Natural transformation assays of the indicated strains using linear tDNA that integrates onto the genome. (**D**) Natural transformation assays of the indicated strains using a replicating plasmid as the tDNA. Data in **C** and **D** are from at least 3 independent biological replicates and shown as the mean ± SD. Statistical comparisons were made by one-way ANOVA with Tukey’s multiple comparison test of the log-transformed data. NS, not significant. *** = *p* < 0.001. LOD, limit of detection. Statistical identifiers directly above bars represent comparisons to the parent.

To test whether the interaction between these proteins is functionally relevant, we mutated 4 residues that lie at the predicted DprA_Vc_-ComM_Vc_ interface (**Fig. 2A**). Two of these residues represented a potential electrostatic interaction (*comM_Vc_ ^R497^*, *dprA_Vc_ ^E353^*), so we mutated these residues by “flipping” their charge: *comM_Vc_ ^R497E^* and *dprA_Vc_ ^E353R^*. We also mutated two leucine residues buried within the interface (*comM_Vc_ ^L455^*, *dprA_Vc_ ^L350^*) to a charged amino acid: *comM_Vc_ ^L455E^*, *dprA_Vc_ ^L350E^*. All of these alleles disrupted DprA_Vc_-ComM_Vc_ interactions. Three alleles – D*prA_Vc_ ^E353R^, omM_Vc_ ^L455E^*, *DprA_Vc_ ^L350E^* – reduced interactions to the limit of detection (**Fig. 2B, S2**), while ComM_Vc_ ^R497E^ exhibited a reduction in interaction (**Fig. 2B, S2**). Consistent with DprA_Vc_-ComM_Vc_ interactions playing a functional role in *V. cholerae*, the 3 alleles that completely eliminated interactions (d*prA_Vc_ ^E353R^, omM_Vc_ ^L455E^*, *dprA_Vc_ ^L350E^*) all phenocopied Δ*comM* in NT frequency assays, while the allele that only reduced interactions (*comM_Vc_ ^R497E^*) had no effect on NT (**Fig. 2C**). Importantly, the point mutations in *dprA* (d*prA_Vc_ ^E353R^* and *dprA_Vc_ ^L350E^*), did not completely inhibit NT (as in Δ*dprA*), indicating that some functions of DprA remain intact (**Fig. 2C**). Furthermore, d*prA_Vc_ ^E353R^* and *dprA_Vc_ ^L350E^* were both epistatic with Δ*comM_Vc_* (**Fig. 2C**).

Together, this suggests that disrupting DprA_Vc_-ComM_Vc_ interactions disrupts ComM_Vc_ activity during NT, which is consistent with what was observed above in *H. pylori* (**Fig. 1**). Also, a multiple sequence alignment of DprA homologs indicated that DprA_Hp_ ^L252^, which is critical for DprA-ComM interactions in *H. pylori* (**Fig. 1B, S2**), is widely conserved in DprA homologs containing a C-terminal winged-helix domain, including in *V. cholerae* DprA_Vc_ ^L354^ (**Fig. S3**). Consistent with a conserved role for this residue, we found that DprA_Vc_ ^L354E^ diminished NT to the level of Δ*comM_Vc_*, and that this mutation is epistatic with Δ*comM_Vc_* (**Fig. 2C**).

To more rigorously test this model, we took advantage of the potential electrostatic interaction between ComM_Vc_ ^R497^-DprA_Vc_ ^E353^. Our results demonstrate that flipping the charge on DprA (*dprA_Vc_ ^E353R^*) disrupts DprA_Vc_ - ComM_Vc_ interactions and ComM activity. Quantitative BACTH assays demonstrate that *comM_Vc_ ^R497E^* also partially disrupts DprA_Vc_-ComM_Vc_ interactions. Based on the AF3 model, the effect of these mutations may be due to electrostatic repulsion between DprA_Vc_ ^E353R^ and ComM_Vc_ ^R497^, with mutation of the DprA side of this interface exhibiting a stronger phenotype. Consistent with an electrostatic interaction, we found that mutating *dprA_Vc_ ^E353^* to lysine (*dprA_Vc_ ^E353K^*), another positively charged amino acid, phenocopied *dprA_Vc_ ^E353R^*, while mutating this residue to glutamine (*dprA_Vc_ ^E353Q^*), an uncharged but similarly sized residue, did not (**Fig. 2B**). If ComM_Vc_ ^R497^-DprA_Vc_ ^E353^ mediate an electrostatic interaction, we hypothesized that flipping the charge on both sides of the interface (*e.g.*, combining *comM_Vc_ ^R497 E^* and *dprA_Vc_ ^E353R^* mutations) should restore ComM_Vc_ activity by restoring DprA_Vc_-ComM_Vc_ interactions. Indeed, we found that NT in *dprA_Vc_ ^E353R^ comM_Vc_ ^R497E^* was partially restored when compared to the *dprA_Vc_ ^E353R^* single mutant (**Fig. 2C, S4**). Importantly, NT of *dprA_Vc_ ^L350E^ comM_Vc_ ^R497E^* was not restored when compared to *dprA_Vc_ ^L350E^* (**Fig. S4**). This indicates that there is specificity to the recovery observed in *dprA_Vc_ ^E353R^ comM_Vc_ ^R497E^*, and demonstrates that *comM_Vc_ ^R497E^* is not simply a gain-of-function allele of ComM. All together, the data strongly suggest that DprA_Vc_ ^E353^ and ComM_Vc_ ^R497^ interact, which further validates the AF3 model.

While we see partial recovery for NT in *dprA_Vc_ ^E353R^ comM_Vc_ ^R497E^*, we did not see recovery of interaction with these proteins in BACTH assays (**Fig. 2B, S2**). One possibility is that the recovered protein-protein interaction is below the detection limit for this assay. This is supported, in part, by the fact that NT is only partially restored in *dprA_Vc_ ^E353R^ comM_Vc_ ^R497E^*.

Prior work shows that ComM is not required for the uptake of tDNA into the cytoplasm, instead, it is only required for the recombination of tDNA into the genome^11,18^. Consistent with this, NT of Δ*comM* is indistinguishable from the parent when a replicating plasmid is used as the tDNA^11^, which does not require recombination (**Fig. 2D**). DprA, however, has a presynaptic role in protecting tDNA from being degraded in the cytoplasm prior to tDNA integration^6^. Consistent with that, NT of Δ*dprA* is significantly reduced when a replicating plasmid is used as tDNA (**Fig. 2D**). The point mutations in DprA_Vc_ that disrupt DprA_Vc_-ComM_Vc_ interactions (*dprA_Vc_ ^L350E^* and *dprA_Vc_ ^E353R^*), however, do not affect NT with a replicating plasmid. This suggests that these DprA alleles retain their presynaptic function; namely, the ability to protect tDNA in the cytoplasm.

### DprAVc-ComMVc interactions are required for branch migration during NT

Our data thus far suggest that DprA_Vc_-ComM_Vc_ interactions are critical for ComM_Vc_ activity because mutations that disrupt this interaction phenocopy a Δ*comM_Vc_* mutant. Also, these mutations are epistatic with a Δ*comM_Vc_* mutation. ComM is required for recombinational branch migration during NT. Thus, we next sought to test whether DprA_Vc_-ComM_Vc_ interactions are critical for ComM_Vc_ -dependent branch migration. To do this, we used a previously established assay to evaluate branch migration activity *in vivo*^11^. This assay indirectly measures branch migration by assessing the comigration of two linked genetic markers: a Trimethoprim antibiotic resistance marker (Tm^R^) and a premature stop codon in *lacZ* (*lacZ*\*) (**Fig. 3A**). We employed two different tDNA products, one where Tm^R^ and *lacZ*\* are separated by 243 bp and another where they are separated by 820 bp. Consistent with ComM_Vc_ playing an important role in branch migration during NT, Δ*comM_Vc_* exhibits significantly reduced comigration for these markers with both tDNA products (**Fig. 3B**). In this assay, we found that the mutations that disrupt DprA_Vc_-ComM_Vc_ interactions (*dprA_Vc_ ^L350E^*, *dprA_Vc_ ^E353R^,* and *comM_Vc_ ^L455E^*) had a significant reduction in comigration that was similar to Δ*comM_Vc_* (**Fig. 3B**).

**Fig. 3.**
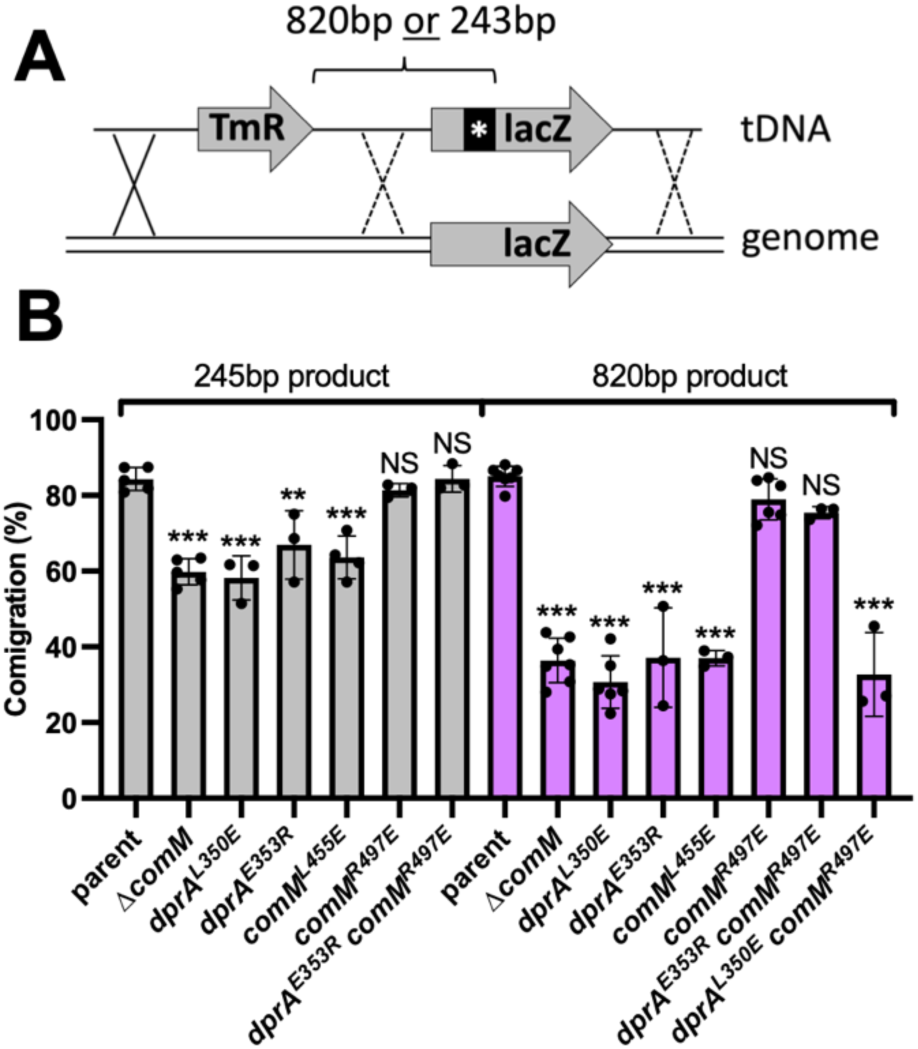
ComMVc-DprAVc interactions are required for branch migration during NT in *V. cholerae*. (**A**) Schematic of the experimental setup to assess branch migration via linkage analysis between a trimethoprim resistance marker (Tm^R^) and a nonsense mutation in *lacZ*. Two different tDNAs were used: one where the *lacZ* nonsense mutation was 243 bp away from the Tm^R^ cassette, and another where the spacing was 820 bp. (**B**) The two tDNA products described in **A** were transformed into the indicated strains and the comigration frequency (i.e., the linkage frequency between the Tm^R^ and *lacZ* nonsense mutation) is reported. All strains in these assays contained a Δ*mutS* mutation to prevent mismatch repair from confounding the results. Data are from at least 3 independent biological replicates and shown as the mean ± SD. Statistical comparisons were made by one-way ANOVA with Tukey’s multiple comparison test of the log-transformed data for each tDNA product. NS, not significant. *** = *p* < 0.001, ** = *p* < 0.01. Statistical identifiers directly above bars represent comparisons to the parent.

Furthermore, we found that comigration in *dprA_Vc_ ^E353R^ comM_Vc_ ^R497E^* was restored when compared to the *dprA_Vc_ ^E353R^* single mutant (**Fig. 3B**). In fact, comigration of *dprA_Vc_ ^E353R^ comM_Vc_ ^R497E^* was restored to parental levels, despite the fact that NT is only partially restored in this background (**Fig. 3B**, **Fig. 2C**). This is consistent with DprA_Vc_-ComM_Vc_ interactions being restored when the charge on both sides of this electrostatic interaction are flipped, which further supports the validity of the AF3 model. Importantly, comigration was not restored in *dprA_Vc_ ^L350E^ comM_Vc_ ^R497E^* when compared to *dprA_Vc_ ^L350E^* indicating that the recovery observed was specific and not due to *comM_Vc_ ^R497E^* being a gain-of-function ComM allele (**Fig. 3B**).

### DprAVc recruits ComMVc to the recombination site during NT

DprA has predominantly been characterized for the presynaptic role it plays during NT. In particular, it plays an important role in recruiting RecA to the tDNA via a direct protein-protein interaction^6–10^. Our data suggest that DprA-ComM interactions are critical for ComM-dependent branch migration. And while it is clear that ComM activity is required for NT and not for other forms of recombination, the mechanism by which this specificity is determined is unclear. Thus, we hypothesized that DprA recruits ComM to the recombination site during NT.

We have previously shown that in single cells undergoing NT, a fluorescent fusion of ComM (GFP-ComM_Vc_) forms foci that are (1) dependent on RecA, (2) only formed in the presence of exogenous tDNA, (3) specifically formed at the chromosomal locus homologous to the tDNA, and (4) transient and highly dynamic, which all together indicates that these foci represent active ComM complexes acting at the recombination site during NT^17^. So, to test whether DprA_Vc_-ComM_Vc_ interactions help recruit ComM_Vc_ to the recombination site, we assessed GFP-ComM_Vc_ focus formation in cells incubated with tDNA (**Fig. 4A**). Consistent with prior work, we found that the parent strain forms many GFP-ComM_Vc_ foci (**Fig. 4B-C**), with ∼60% of cells exhibiting at least 1 GFP-ComM_Vc_ focus within the timelapse imaging window. By contrast, point mutations that eliminate DprA_Vc_ - ComM_Vc_ interactions (*dprA_Vc_ ^L350E^*, *dprA_Vc_ ^E353R^*, and *comM_Vc_ ^L455E^*) reduce ComM_Vc_ focus formation to the limit of detection (**Fig. 4B-C**). Consistent with a partial reduction in DprA_Vc_-ComM_Vc_ ^R497E^ interactions by BACTH (**Fig. S2**), *comM_Vc_ ^R497E^* reduces ComM_Vc_ focus formation compared to the parent (**Fig. 4B-C**). Also, consistent with a reduction in DprA_Vc_ ^E353R^-ComM_Vc_ ^R497E^ interactions by BACTH (**Fig. 2B, S2**), *dprA_Vc_ ^E353R^ comM_Vc_ ^R497E^* reduces ComM_Vc_ focus formation almost to the limit of detection (**Fig. 4B-C**). Importantly, GFP-ComM_Vc_ expression is not altered in any of the mutants tested (**Fig. S5**), indicating that disruption of DprA_Vc_-ComM_Vc_ interactions does not alter ComM_Vc_ expression and/or stability.

**Fig. 4.**
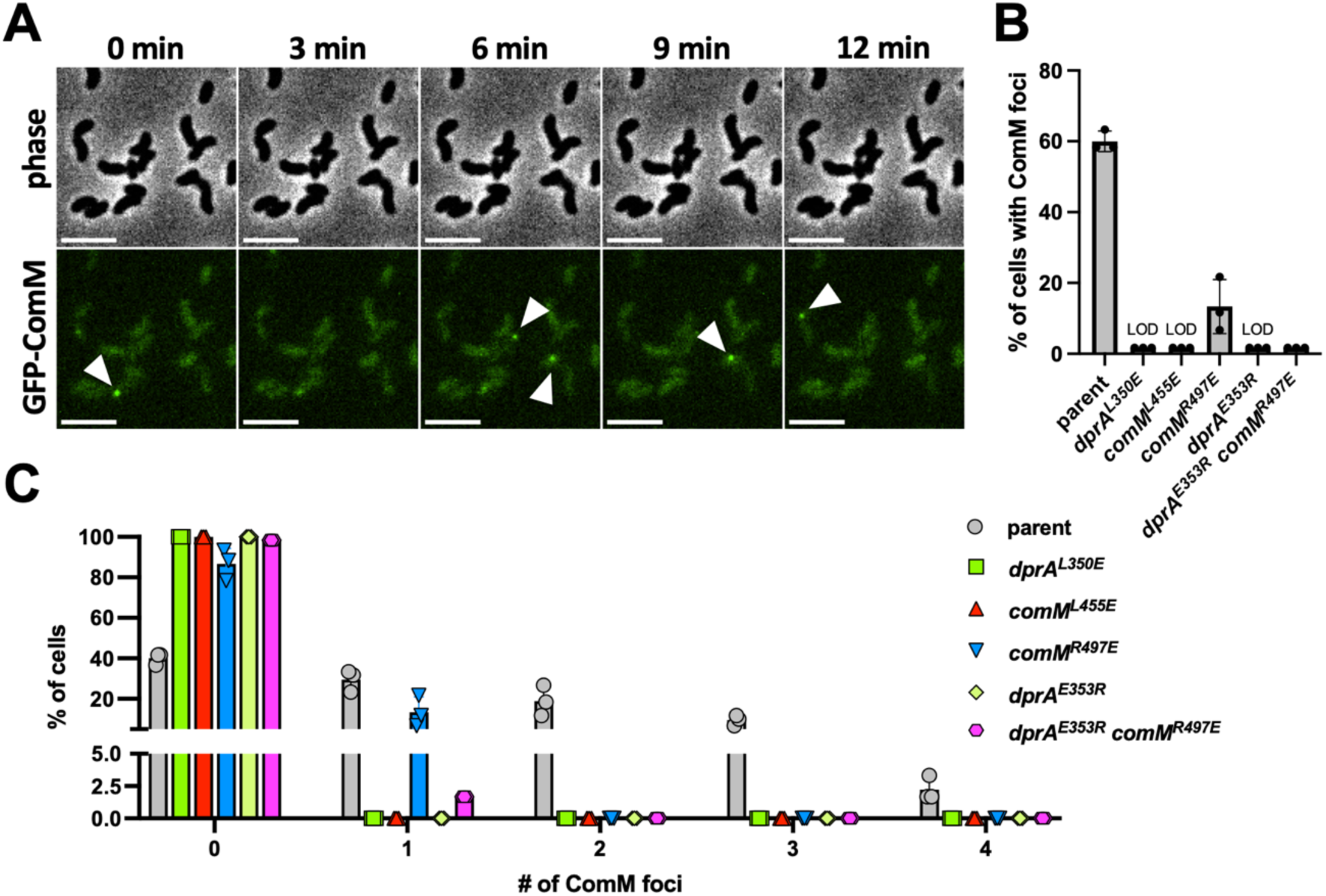
DprAVc recruits ComMVc to the recombination site during NT in *V. cholerae*. (**A**) Montage of timelapse imaging showing that the recruitment of ComM_Vc_ to tDNA can be easily observed as fluorescent foci (white arrows) in a parent strain harboring a functional GFP-ComM_Vc_ fusion at the native locus. Scale bar, 4µm. Quantification of timelapse images for (**B**) the percent of cells that exhibited at least one GFP-ComM_Vc_ focus and (**C**) the distribution of the number of GFP-ComM_Vc_ foci observed per cell. The indicated strains were imaged every 3 min for 1.5 hours. For each replicate (*n* = 3), 60 cells were analyzed and the data are shown as the mean ± SD. LOD, limit of detection.

Interestingly, the reduction in ComM_Vc_ ^R497E^ focus formation does not equate to a reduction in NT frequency for *comM_Vc_ ^R497E^* (**Fig. 2C**), which suggests that the GFP-ComM_Vc_ focus formation assay can reveal more subtle defects in recombination compared to NT frequency assays. Also, we show above that the branch migration deficit of *dprA_Vc_ ^E353R^* is completely restored in *dprA_Vc_ ^E353R^ comM_Vc_ ^R497E^* (**Fig. 3B**). Despite this, we see almost no restoration in GFP-ComM_Vc_ focus formation in *dprA_Vc_ ^E353R^ comM_Vc_ ^R497E^* (**Fig. 4C**; exactly one GFP-ComM focus was observed in the 60 cells analyzes in each replicate). This is perhaps not surprising, however, because *dprA_Vc_ ^E353R^ comM_Vc_ ^R497E^* only partially rescues the transformation deficit of *dprA_Vc_ ^E353R^*. Thus, for mutations that completely disrupt DprA_Vc_-ComM_Vc_ interactions (*dprA_Vc_ ^L350E^*, *dprA_Vc_ ^E353R^*, and *comM_Vc_ ^L455E^*) we find that these mutants phenocopy Δ*comM* in both NT frequency and branch migration assays. For mutations that only partially disrupt DprA_Vc_-ComM_Vc_ interactions (*comM_Vc_ ^R497E^* and *dprA_Vc_ ^E353R^ comM_Vc_ ^R497E^*), however, we observe only a partial restoration in NT frequency, but a complete restoration in branch migration.

Together, these data are consistent with a model in which DprA recruits ComM to the recombination site during NT. If these interactions are partially disrupted, this diminishes the frequency of recombination events that proceed to ComM-dependent branch migration (explaining the observed reduction in NT frequency and GFP-ComM focus formation in *comM_Vc_ ^R497E^* and *dprA_Vc_ ^E353R^ comM_Vc_ ^R497E^*); but for the recombination events where ComM is successfully recruited in these backgrounds, branch migration is highly effective (explaining the parent levels of branch migration observed in *comM_Vc_ ^R497E^* and *dprA_Vc_ ^E353R^ comM_Vc_ ^R497E^*).

### DprA promotes ComM-dependent branch migration for NT in species-specific manner

We show above that DprA-ComM interactions are preserved in both *H. pylori* and *V. cholerae*. Also, AF3 analysis indicates that DprA-ComM interactions are likely also broadly conserved in species that contain a DprA homolog with a C-terminal winged-helix domain (**Dataset S1**). Despite this potentially broad conservation in DprA-ComM interactions, the interaction interfaces between diverse homologs may be distinct. For example, the electrostatic interaction that we demonstrate is critical for DprA_Vc_-ComM_Vc_ interactions, is absent in the DprA_Hp_-ComM_Hp_ model (**Fig. 1A, 2A**), and is not conserved at the sequence level (**Fig. S6**). Consistently, ectopically expressing ComM_Hp_ in *V. cholerae* does not complement the defect in NT and branch migration of *V. cholerae* Δ*comM* mutant (**Fig. 5A-B**). Thus, we hypothesized that DprA-ComM interactions may be species specific (*i.e.*, DprA_Vc_ cannot recruit ComM_Hp_ and *vice versa*). If this is the case, ectopic expression of both DprA_Hp_ and ComM_Hp_ in *V. cholerae* should be sufficient to rescue ComM activity. First, we find that ectopic expression of DprA_Hp_ in *V. cholerae* results in a slight decrease in NT frequency, but deletion of native *comM* in this background still diminishes NT ∼100-fold as expected (**Fig. 5**). When DprA_Hp_ and ComM_Hp_ are both ectopically expressed in *V. cholerae*, we found that NT was unchanged when native *comM* is deleted in this background (**Fig. 5A**) and branch migration activity was restored (**Fig. 5B**). This result is consistent with DprA_Hp_ recruiting ComM_Hp_ to facilitate ComM-dependent branch migration activity in *V. cholerae*. These results also suggest that DprA and ComM represent the “functional unit” that facilitates branch migration during NT in Gram-negative species.

**Fig. 5.**
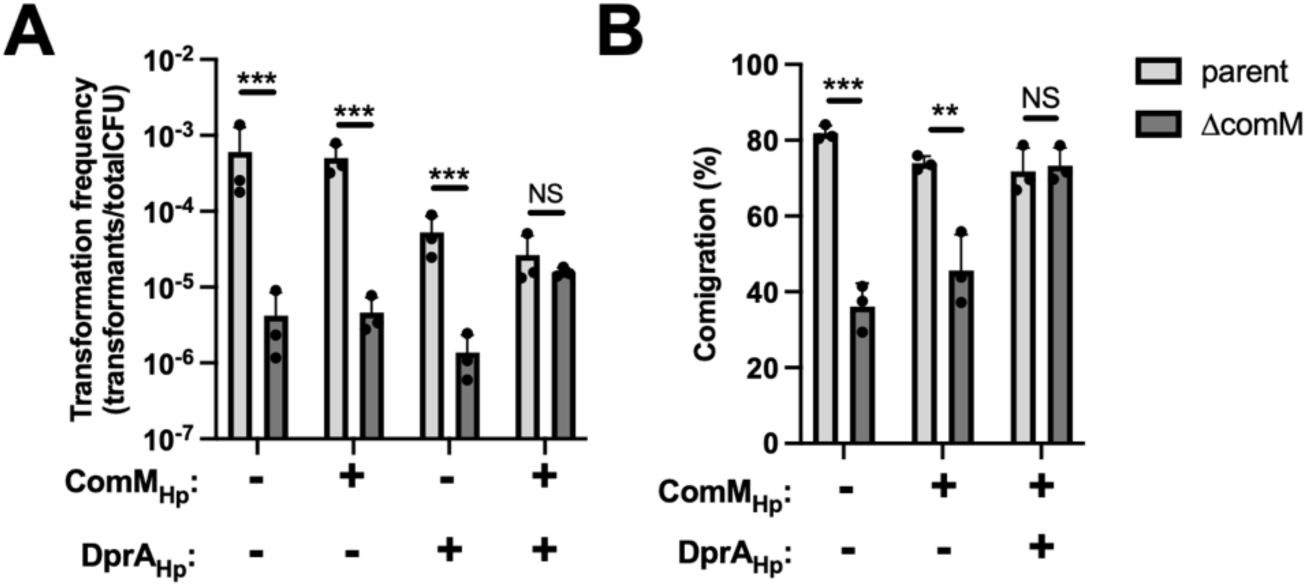
DprA-dependent recruitment of ComM is species-specific. (**A**) Transformation frequency of *V. cholerae* strains where native *comM* is intact (parent; gray bars) or deleted (Δ*comM*; black bars) using linear tDNA that integrates onto the genome. Strains express *H. pylori* ComM (ComM_Hp_) and/or DprA (DprA_Hp_) from an ectopic locus as indicated (+ denotes that the strain expresses the indicated protein). (**B**) Comigration frequency in the indicated strain backgrounds to assess branch migration via linkage analysis between a trimethoprim resistance marker (Tm^R^) and a nonsense mutation in *lacZ* as depicted in Fig. 3A. For the tDNA used in this experiment the *lacZ* nonsense mutation was 820 bp away from the Tm^R^ cassette. All data are from at least 3 independent biological replicates and shown as the mean ± SD. Statistical comparisons were made by one-way ANOVA with Tukey’s multiple comparison test of the log-transformed data. NS, not significant. *** = *p* < 0.001, ** = *p* < 0.01.

## DISCUSSION

The results presented above advance our understanding of the molecular coordination underlying tDNA integration during horizontal gene transfer by NT. Specifically, they demonstrate that in *H. pylori* and *V. cholerae*, the winged-helix domain of DprA directly binds to ComM to recruit this branch migration factor to the recombination site during NT. Our bioinformatic analysis also suggests that DprA-ComM interactions are conserved in other Gram-negatives. Thus, it is tempting to speculate that this activity is broadly conserved in other naturally transformable Gram-negative species.

The structure of *Legionella* ComM was recently solved by the Fronzes group^19^. Using AF3, we modeled a ComM_Vc_ hexamer with DNA, both without and with DprA_Vc_. We confirmed that the AF3 model of the ComM_Vc_ hexamer with DNA aligns very well to the solved structure (**Fig. S7**). The DprA^C-ter^_Vc_-ComM_Vc_ interface used in our work superimposes perfectly with the larger AF3 model including a ComM_Vc_ hexamer, DprA_Vc_ and DNA. The latter model suggests that the DprA-ComM interaction interface does not conflict with the ComM interfaces required for hexamerization or DNA-binding (**Fig. S8**). Interestingly, the DprA-ComM interface involves the ComM Mg chelatase domain^19^. The function of this domain in ComM is poorly understood, but our results suggest that one of its functions is to aid in recruitment of this helicase during NT.

Work from our group and others has characterized the molecular and structural basis of ComM-dependent branch migration of D-loops during NT in Gram-negative species^11,12,19^. D-loop structures are also generated as intermediates during DNA repair, however, ComM does not facilitate this process and is instead uniquely required for tDNA integration during NT^11^. This suggests that ComM may be specifically loaded onto the D-loops generated during NT. However, the basis for ComM loading has remained elusive. Our results demonstrate DprA is likely required to load ComM onto D-loops during NT. Thus, DprA-dependent loading of ComM may explain why this branch migration factor is exclusively associated with recombination during NT.

Our prior work has demonstrated that ComM is a bidirectional helicase that can translocate both 5’ → 3’ and 3’ → 5’ on model branched DNA substrates *in vitro*^11^. If this directionality is unregulated, then ComM would facilitate (*i.e.*, expand the D-loop) and inhibit (*i.e.*, collapse the D-loop) tDNA integration with equal probability. Indeed, this is exactly what is observed *in vitro* with purified ComM^11^. Single cell tracking of the spatiotemporal dynamics of tDNA integration, however, suggest that the recombination reaction is highly efficient^17^, which suggests that there may be additional factors that regulate ComM directionality *in vivo*. A recent study highlights that ComM works in conjunction with a NT-specific nuclease, YraN, which processes D-loops and may facilitate directional branch migration by cleaving the displaced genomic strand^12^. Here, we show that DprA-ComM interactions are required for branch migration activity *in vivo*. Thus, it is tempting to speculate that DprA-dependent loading of ComM onto the D-loop also helps regulate the directionality of branch migration, which is something we will test moving forward.

DprA-dependent recruitment of ComM is notable for a number of reasons. DprA has primarily been characterized for its presynaptic roles during NT. This includes its role in protecting single-stranded tDNA in the cytoplasm, eviction of SSB, and recruitment of RecA^6–10^. Together, these activities help promote RecA-mediated strand invasion to initiate the recombination process. Our results demonstrate that DprA is also required to initiate branch migration, which indicates that it also plays a postsynaptic role during NT by recruiting ComM. DprA has also been implicated in playing a regulatory role in some systems like *Streptococcus pneumoniae*, where DprA directly interacts with the master regulator of competence, ComE, to shut-off competence^20^. Due to its varied roles during NT, most of which are facilitated by direct protein-protein interactions, it is tempting to speculate that DprA serves as the master “hub” that coordinates this complex process. On a related note, ComM has previously only been implicated in the postsynaptic step of branch migration. The recruitment of ComM by DprA may also suggest that ComM aids in a presynaptic step of recombination (*e.g.*, the homology search or initiation of strand invasion).

The highly varied roles for DprA during NT also highlight the value of the approach used in this study. Because of the critical presynaptic roles that DprA performs during NT, deletion of DprA completely inhibits NT. Thus, uncovering its postsynaptic role would not have been possible using classical genetic approaches like transposon screens. While genetic screens can help identify the genes and proteins necessary for a particular phenotype, they may fail to reveal the underlying molecular coordination necessary for their activity – especially in cases where factors have more than one function. The structural modeling approach used here, however, helped reveal unexpected interactions that led to testable hypotheses, which ultimately uncovered how these factors coordinate to carry out their activity. Advances in structural modeling are developing at breakneck speed, and recent work demonstrates that genome-wide unbiased searches for novel protein-protein interactions are now highly feasible^21,22^. These unbiased genome-wide *in silico* screens hold great promise in accelerating the discovery of these types of unexpected interactions moving forward.

## METHODS

### Bacterial strains and culture conditions

All *V. cholerae* strains used throughout this study are derivatives of strain E7946^23^. Cells were routinely grown in LB Miller broth and agar supplemented with spectinomycin (200 µg/mL), kanamycin (50 µg/mL), trimethoprim (10 µg/mL), tetracycline (0.5 µg/mL), sulfamethoxazole (100 µg/mL), chloramphenicol (1 µg/mL), erythromycin (10 µg/mL), carbenicillin (20 µg/mL), and/or zeocin (100 µg/mL) as appropriate.

All *H. pylori* strains used in this study are derivatives of strain 26695^24^. Cultures were grown under microaerobic conditions (5% O_2_, 10% CO_2_, using the Don Whitley system) at 37°C. Blood agar base medium (BAB) supplemented with 10% defibrillated horse blood (ThermoFisher Scientific) was used for plate cultures. An antibiotic mix containing polymyxin B (0.155 mg/ml), vancomycin (6.25 mg/ml), trimethoprim (3.125 mg/ml), and amphotericin B (1.25 mg/ml) was added to both plate and liquid cultures. Additional antibiotics were added as required: kanamycin (20 μg/ml), apramycin (12.5 μg/ml), and chloramphenicol (8 μg/ml) streptomycin (10 μg/ml).

### Construction of mutant strains

For *V. cholerae*, all strains were generated by natural transformation using splicing-by-overlap-extension (SOE) PCR mutant constructs and cotransformation exactly as previously described^25,26^. Ectopic expression constructs were generated by SOE PCR and integrated onto the *V. cholerae* genome exactly as previously described^27^. Unless otherwise stated, mutations to *dprA_Vc_* and *comM_Vc_* were made at the native locus. The *gfp-comM_Vc_* translational fusion was generated at the native locus as previously described^28^.

For generating mutant variants in *H. pylori* the specific gene regions (with 200bp of flanking sequences) were amplified from genomic DNA (gDNA) of *H. pylori* 26695 using sequence specific primers and cloned into pjET1.2 plasmid using sequence- and ligation-independent cloning (SLIC)^29^. Replacement of the ORF by antibiotic resistant cassettes (either chloramphenicol, kanamycin or apramycin) was carried out by SLIC. The constructs were introduced in *H. pylori* by natural transformation and at least two independent clones were selected for each construct. For complementation by expression from the *ureA* locus the corresponding gene ORFs were amplified by PCR and cloned by SLIC into a pjET2.1 plasmid carrying an antibiotic resistance cassette flanked by 200 bp upstream and downstream of the *ureA* gene. The resulting construct was introduced in *H. pylori* by natural transformation. Strain *ΔdprA::Kan^R^, P_ureA_ ::dprA^ΔC-ter^::Apra^R^* expressing a DprA protein lacking residues 210-270 was previously described^13^.

All strains were confirmed by PCR and/or sequencing. For a detailed list of all strains, see **Table S1.** For a complete list of oligos used to generate mutant constructs, see **Table S2**.

### Bioinformatic analysis of DprA homologs

Using BLAST^30^, we searched for DprA homologs in a set of bacterial genomes where ComM homologs had previously been searched^11^. For all identified DprA homologs, we performed protein structure prediction using AlphaFold v2.3^31^ as implemented in the ColabFold v1.5.2 pipeline^32^. We ran MMseqs2^33^ against the UniRef30_2202 database to generate multiple sequence alignments used as input to the AlphaFold2 structure prediction module. We then generated 5 structural models for each DprA homolog using 3 recycles. We structurally superimposed the best models (highest pTM score for each species) with PyMOL (v3.0) and aligned them to the experimental structure of the DprA_Hp_ C-terminal domain^13^ (PDB identifier: 6GW7). We visually identified the presence of a C-terminal winged helix domain in all species with an identified ComM homolog, apart from *B. burgdoferi* (**Fig. S1** and **Dataset S1**). To validate the presence of a C-terminal winged helix domain, we annotated Pfam^34^ domains in all DprA homolog sequences by HH-suite homology search^35^: we ran 5 iterations of HHblits against the UniRef30_2202 database to generate multiple sequence alignments, used as input for HHsearch against Pfam v35.0 (with 80% maximum pairwise sequence identity). We identified a winged helix Pfam domain (PF17782, PF10771.12 or PF15977.8) with HHsearch probability above 90% in all species with ComM homologs, except for *E. coli* (probability 88.2%), *F. nucleatum* (probability 89.7%)*, H. pylori* (probability 23%)*, C. jejuni* (probability 31.7%), and *L. interrogans* (probability 59%). Note that the DprA_Hp_ C-terminal domain is not annotated in Pfam despite being structurally characterized.

### Natural transformation assays

For *V. cholerae*, strains were engineered to bypass the native signals required for NT. Specifically, strains had *P_tac_-tfoX* and Δ*luxO* mutations, which ectopically express the master competence regulator (TfoX) and genetically lock cells in a high cell density state as previously described^36^. Chitin-independent transformation assays to determine NT frequency were performed exactly as previously described^36^. Briefly, cells were grown at 30°C rolling in LB supplemented with 100 µM IPTG, 20 mM MgCl_2_, 10 mM CaCl_2_ to late-log phase. Then, ∼10^8^ CFUs were diluted into instant ocean medium (7g/L; Aquarium Systems). Then, 100 ng of transforming DNA (tDNA) was added to each reaction. The tDNA used to assess transformation frequency was a linear ΔVC1807::Erm^R^ PCR product. VC1807 is a frame-shifted transposase gene. Control reactions were also performed for each strain where no tDNA was added. Cells were incubated with tDNA overnight at 30°C static. Then, reactions were outgrown by adding 1mL of LB and shaking at 37°C for 3 hours. Reactions were then plated for quantitative culture by dribble plating on LB agar supplemented with erythromycin to quantify transformants, and plain LB to quantify total viable counts. Transformation frequency is defined as the CFU/mL of transformants divided by the CFU/mL of total viable counts.

For assays where branch migration was indirectly assessed via linkage analysis in *V. cholerae*, transformation reactions were performed exactly as described above except (1) a different tDNA is used and (2) reactions were plated onto LB agar supplemented with trimethoprim and X-gal (40 µg/mL). The tDNA used in these experiments was previously described^37^ and is schematically depicted in **Fig. 3A**. Briefly, this tDNA consists of (in order from 5’→3’): ∼3 kb homology upstream of *lacZ*, a Tm^R^ cassette, a nonsense mutation in *lacZ* that is situated either 243 bp or 820 bp away from the TmR cassette, and ∼0.9 kb of homology downstream of the point mutation. Thus, when plating these reactions on LB+Tm+X-gal, all colonies will have integrated the Tm^R^ marker, and if cells integrated the genetically linked *lacZ* nonsense mutation they would make white colonies. If they did not integrate the linked lacZ nonsense mutation, they would make blue colonies on these plates. Thus, the comigration frequency is defined as the percent of Tm^R^ colonies that were white on these plates.

Natural transformation frequencies for *H. pylori* were determined as follows. Total chromosomal DNA (600 ng) from a streptomycin resistant 26695 strain was mixed with 45 μL of a culture at OD_600_ = 4.0. Then, three 15μL drops of the mix were spotted on solid medium and incubated overnight at 37°C. Next day, cells were resuspended in 300 μL of peptone water and 10 μl drops of serial dilutions were deposited on BAB plates with and without streptomycin (10 μg/mL). Transformation frequency is defined as the CFU/mL of streptomycin resistant transformants divided by the CFU/mL of total viable counts.

### Microscopy for GFP-ComM focus formation

Assays to assess GFP-ComM focus formation in the presence of tDNA were performed exactly as previously described^17^. Briefly, cells were grown as described above for natural transformation assays. Then, ∼10^7^ cells were mixed with ∼650 ng of a linear ∼6 kb tDNA product that is homologous to the host chromosome, and placed under an 0.2% gelzan pad. Samples were then subjected to timelapse imaging. Phase contrast and fluorescence images were collected every 3 min for a total of 90 mins on a Nikon Ti-2 microscope using a Plan Apo 60X objective, a GFP filter cube, a Hamamatsu ORCA Flash 4.0 camera and Nikon NIS Elements imaging software. The number of GFP-ComM foci formed throughout the timelapse was determined by analyzing 60 cells per replicate.

### Bacterial Adenylate Cyclase Two-Hybrid (BACTH) Assays

BACTH analysis was performed essentially as previously described^16^. Gene inserts were amplified by PCR and cloned into the pUT18C, pUT18, pKT25, or pKNT25 vectors (Euromedex) to generate N- and C-terminal fusions to the T18 and T25 fragments of adenylate cyclase. Miniprepped vectors were then cotransformed into *E*. *coli* BTH101 (Euromedex).

For qualitative assessment of interactions, single colonies were picked and grown in LB supplemented with kanamycin 50 μg/mL and carbenicillin 100 μg/mL. Next, 3 μL of each strain was spotted onto LB agar plates supplemented with kanamycin 50 μg/mL, carbenicillin 100μg/mL, 0.5 mM IPTG and 40 μg/mL X-gal. Plates were incubated at 30°C for 24 hours prior to imaging.

To quantify the strength of BACTH interactions for *H. pylori* homologs, five colonies were collected and used to inoculate 3 mL of LB containing both antibiotics and allowed to grow overnight at 30°C. The OD_600_ of the culture was determined and cells from 1mL were pelleted and resuspended in 1mL of Z Buffer (60 mM Na_2_HPO_4_, 40 mM NaH_2_PO_4_, 10 mM KCl, 1 mM MgSO_4_, pH 7) supplemented with 50 mM β-mercaptoethanol and 0.005% SDS. Next, 100 μL of chloroform was added and reactions were vortexed for 20 s. Then, 20 μL of the upper phase was mixed in 96-well microplates with 180 μL of Z Buffer supplemented with 0.7 mg/mL of ortho-nitrophenyl-β-galactoside (ONPG). Absorbance was then measured kinetically at 540 nm and 414 nm every 5 mins for 2 hours on a BMG Labtech CLARIOStar plate reader.

To quantify the strength of BACTH interactions for *V. cholerae* homologs, colonies were grown in LB containing 50 µg/mL kanamycin, 100 µg/mL carbenicillin, and 500 µM IPTG at 30°C rolling overnight. Then, cultures were washed and adjusted to an OD_600_ = 0.5 in 1 mL Z Buffer supplemented with 50 mM β-mercaptoethanol. Next, 50 µL of 1% SDS and 100 µL of chloroform were added and reactions were vortexed for 15 s. Then, reactions were centrifuged and 50 µL of the aqueous phase was transferred to a 96-well plate containing 150 µL of Z Buffer supplemented with 3.33 mg/mL ONPG. Absorbance was kinetically measured at 420 nm and 550 nm every 10 mins for 10 hours on a Biotek H1M plate reader.

β-galactosidase activity (A) was defined as: A = 1000 * (OD_414/420_ (t) – (1.75 * OD_540/550_ (t)) / (t * V_sample_ * initial OD_600_) and expressed as Miller units: A_414 /420_/(min * mL * OD_600_).

### Western Blot Analysis

*V. cholerae* cells were grown exactly as described above for natural transformation assays. Cells were concentrated to an OD_600_ = 100.0. Then, samples were lysed by mixing 1:1 with 2X SDS Sample Buffer (222 mM Tris pH 6.8, 26 % glycerol, 3.9 % SDS, 0.022 % bromophenol blue, 0.715 M beta-mercaptoethanol) and boiling for 10 mins. Next, 5 µL of each lysed sample was electrophoretically separated on a 10% SDS PAGE gel and transferred to an Immobilon-FL PVDF membrane (Millipore). Membranes were then blocked in Tris-buffered saline containing 0.1% Tween 20 (TBST) supplemented with 5 % milk for 1 hour at room temp. Then, membranes were incubated with a rabbit polyclonal anti-GFP primary antibody (ThermoFisher) overnight at room temp in TBST + 2% milk powder. Next, blots were washed in TBST to remove excess primary antibody, and then incubated for 3 hours at room temp in TBST containing an anti-rabbit HRP secondary antibody. Next, blots were washed in TBST to remove excess secondary antibody, and then incubated with an enhanced chemiluminescence substrate (Pierce) for 1 min at room temp prior to imaging for chemiluminescence on a ProteinSimple FluorChem R instrument.

### Multiple Sequence Alignments

MSAs were generated with MAFFT using the E-INS-i protocol^38^. The MSAs were visualized and colored with Jalview^39^.

### Visualization of Protein Models

Images of protein models were prepared in ChimeraX (v1.9)^40^.

### Statistics

Statistical differences were assessed using GraphPad Prism software. The statistical tests employed are indicated in figure legends. Descriptive statistics for all samples, and a comprehensive list of statistical comparisons can be found in **Dataset S2**.

## ACKNOWLEDGEMENTS

We thank Carrie Shaffer and Prashant Damke for providing *H. pylori* genomic DNA to ABD. We also thank Anne Marie Di Guilmi for setting up the initial BACTH experiments and Juliette Bernardin for technical assistance in JPR’s laboratory. This work was supported by grant R35GM128674 from the National Institutes of Health to ABD and grant ANR-22-CE44-0044 from Agence Nationale de la Recherche to JPR, JA and RG. We thank the BIOI2 platform for making the ColabFold pipeline easily accessible at the I2BC for YF, JA and RG.

---

**Fig. S1.**
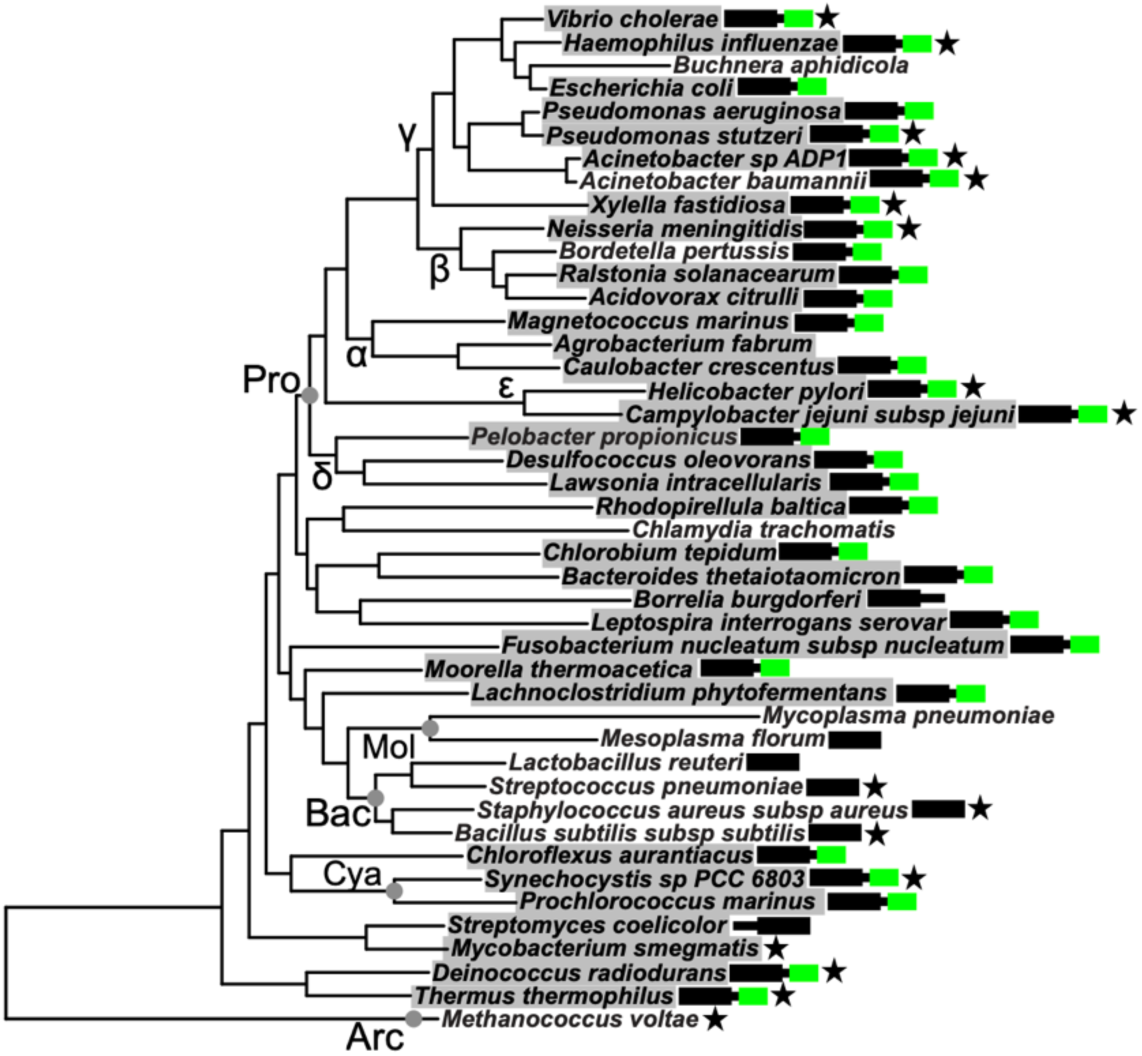
Most species with ComM encode a DprA homolog with a C-terminal winged-helix domain. Estimated maximum likelihood phylogeny of the indicated species based on a concatenated alignment of 36 conserved proteins identified from whole genome sequences (Genome Accession numbers are available in **Dataset S1**). Species that encode a ComM homolog are highlighted in gray. The architecture of the DprA homolog (if present) is indicated to the right of each species name. The core DprA smf domain is indicated by a black box, while the presence of a C-terminal winged helix domain is demarcated by a green box (details of C-terminal domain identification are presented in **Methods** and **Dataset S1**). Species that have been shown to have the capacity for NT are designated by a black star. Major taxa are labeled along their nodes. Pro: Proteobacteria (Greek letters denote subdivisions); Bac: Bacilli; Mol: Mollicutes; Cya: Cyanobacteria; Arc: Archaea.

**Fig. S2.**
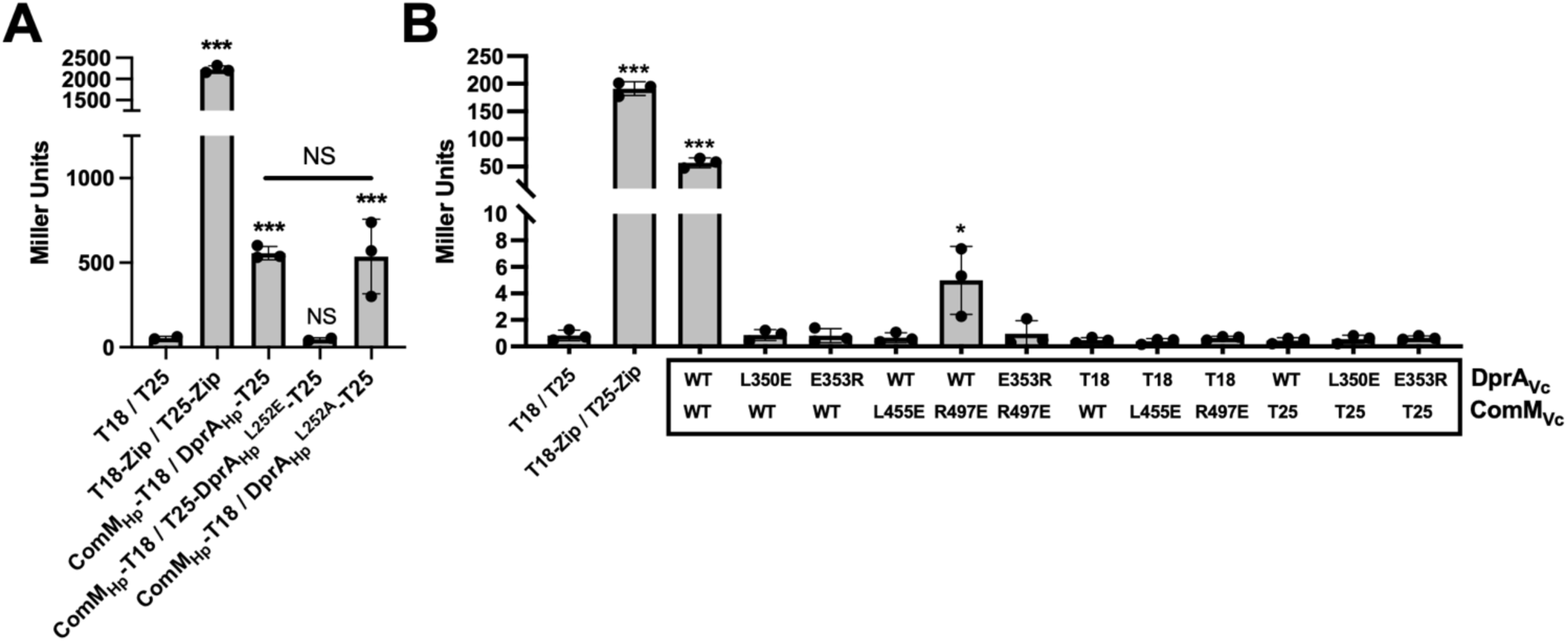
Quantification of BACTH for ComM-DprA interactions. BACTH for the indicated protein pairs in (**A**) *H. pylori* and (**B**) *V. cholerae* were quantitatively assessed for interactions by performing Miller Assays. Statistical comparisons were made by one-way ANOVA with Tukey’s multiple comparison test of the log-transformed data. Statistical identifiers directly above bars represent comparisons to the parent containing the empty vectors (T18 / T25). NS, not significant. *** = *p* < 0.001.

**Fig S3.**
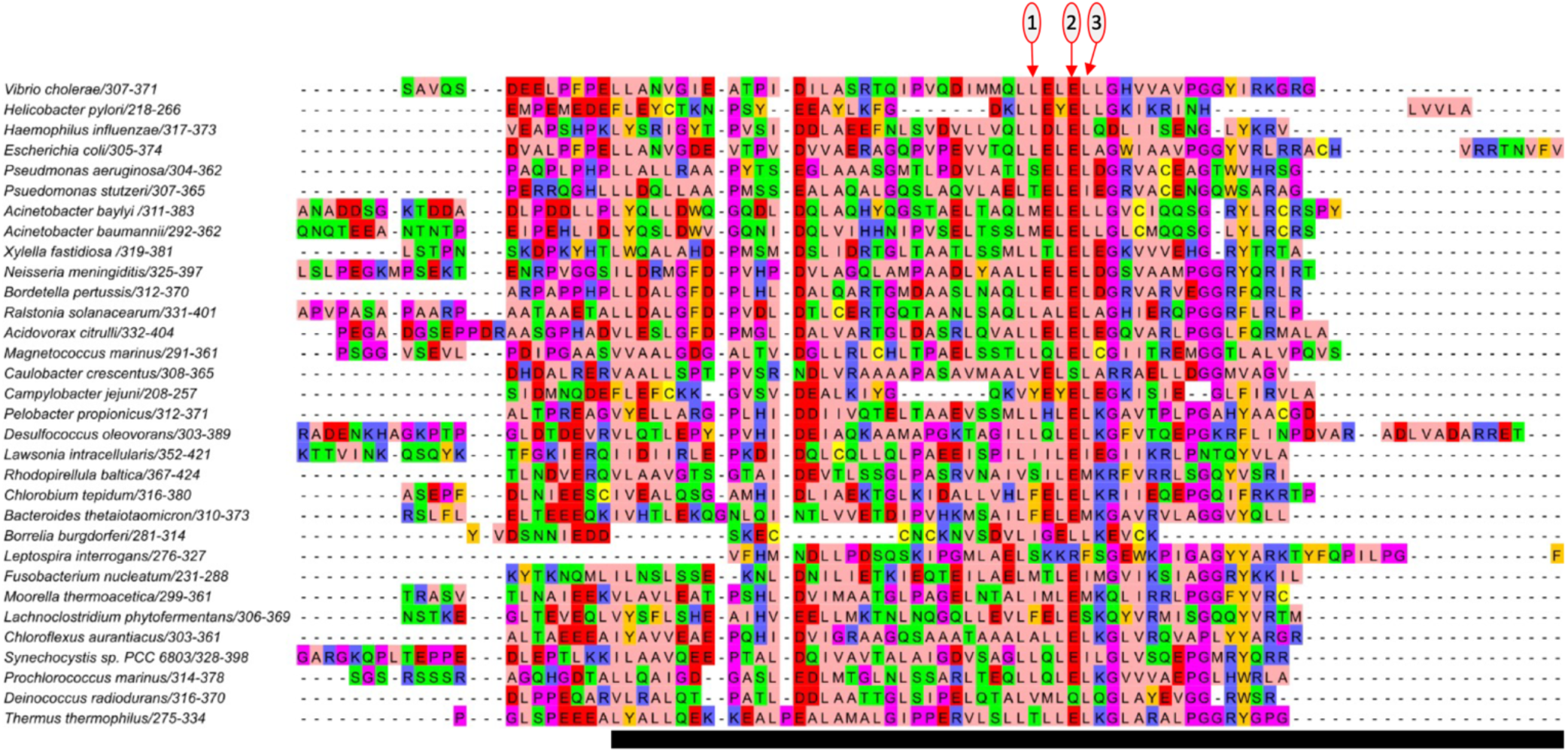
MSA of the C-terminus of DprA homologs with a winged helix domain. MSA of DprA homologs from species shown in **Fig S1** encoding a DprA homolog including a C-terminal winged helix domain. For easier pairwise comparison, the *Helicobacter pylori* sequence was shifted to just below the *Vibrio cholerae* sequence. The MSA is focused on the C-terminal winged helix domain. Amino acids are colored following their physicochemical properties based on the Zappo color scheme. Numbered positions denote: (1) DprA_Vc_ ^L350^ and DprA_Hp_ ^L248^; (2) DprA_Vc_ ^E353^; and (3) DprA_Vc_ ^L354^ and DprA_Hp_ ^L252^. The black bar below sequences denotes the portion of each DprA homolog used for AF3 analysis in **Dataset S1**.

**Fig. S4.**
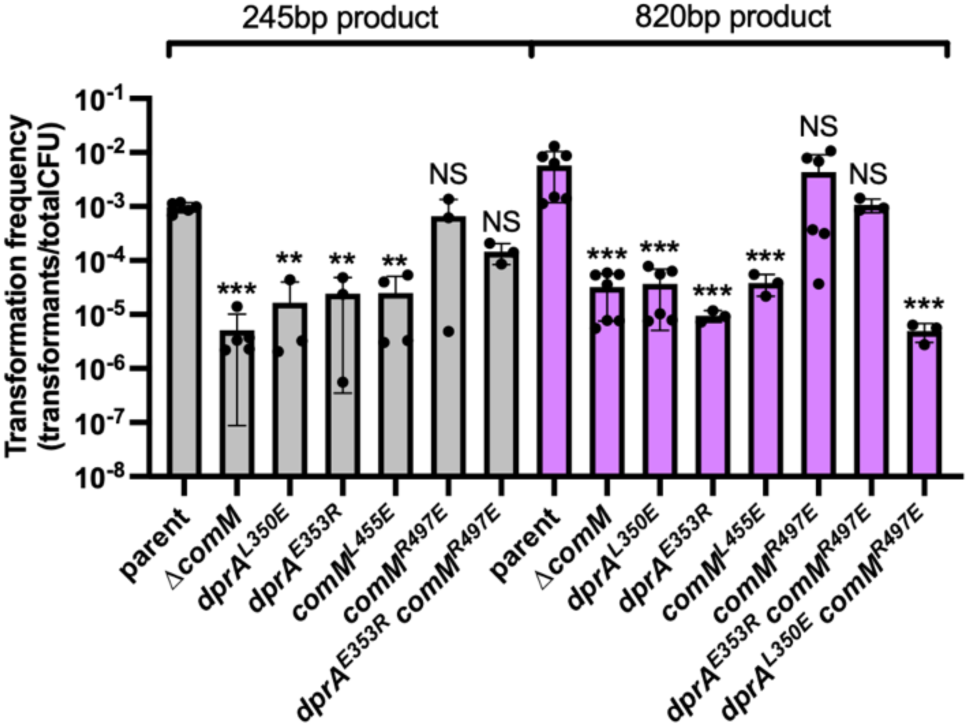
NT frequencies for branch migration assays. NT frequency data for the branch migration assays shown in Fig. 3B. Data are from at least 3 independent biological replicates. Statistical comparisons were made by one-way ANOVA with Tukey’s multiple comparison test of the log-transformed data. NS, not significant. *** = *p* < 0.001, * = *p* < 0.05. Statistical identifiers directly above bars represent comparisons to the parent.

**Fig. S5.**
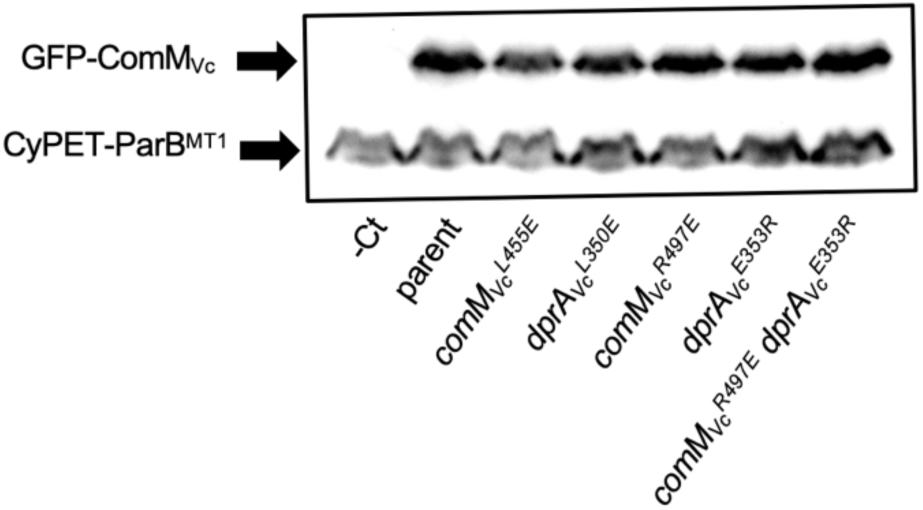
ComMVc protein levels is not affected when ComMVc-DprAVc interactions are disrupted. Western blot analysis of the indicated strains using *α*-GFP antibodies. All strains contained a *P_lac_-cyPET-parB^MT1^* construct, the product of which (∼61 kDa) cross-reacts with *α*-GFP antibodies and serves as a loading control. The –Ct strain lacked a GFP-ComM construct. While all remaining strains harbored a GFP-ComM (∼82 kDa) construct at the native locus along with the indicated point mutations.

**Fig. S6.**
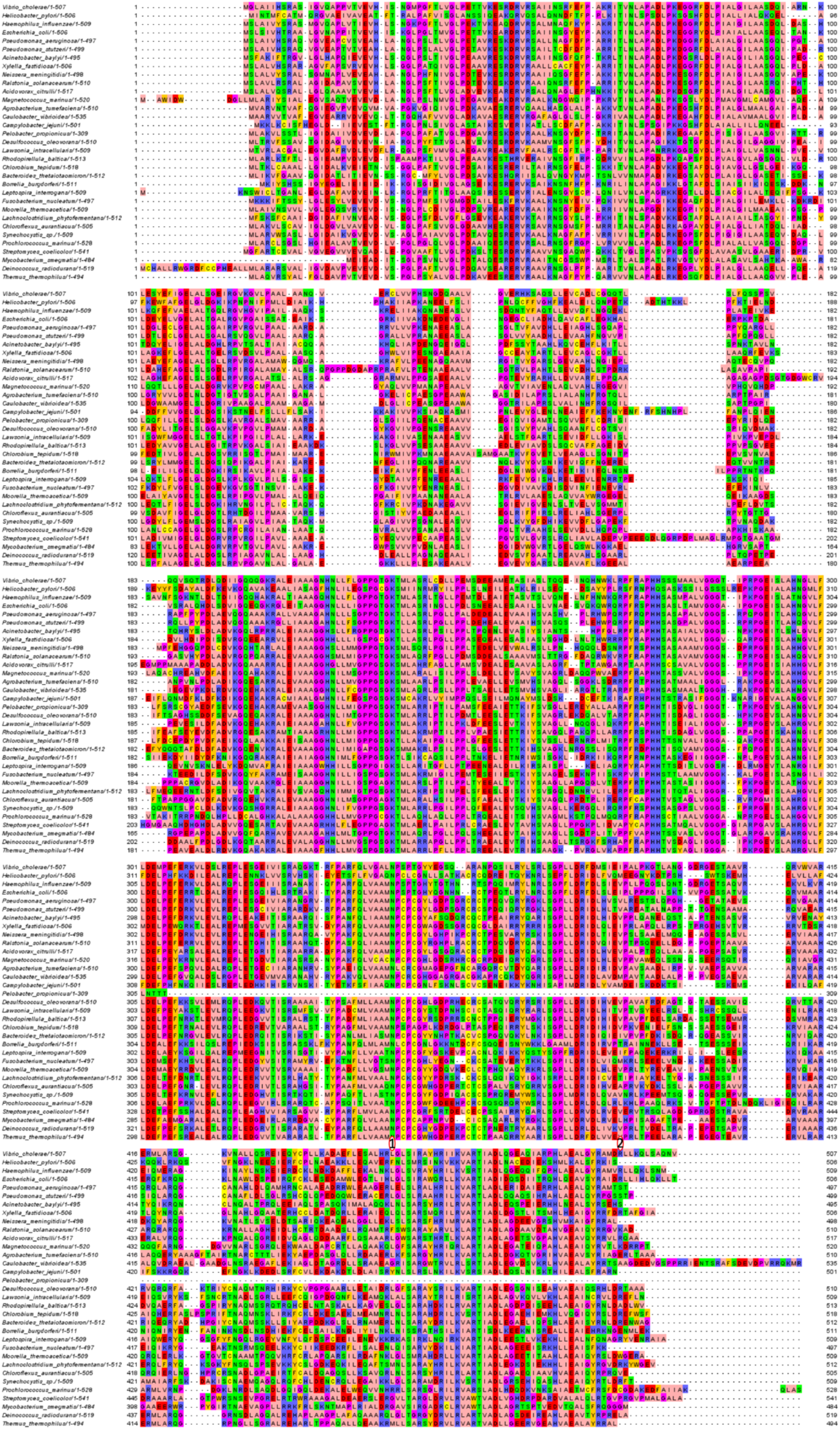
MSA of ComM homologs. MSA of ComM homologs from species shown in **Fig S1**. Amino acids are colored following their physicochemical properties based on the Zappo color scheme. Numbered positions denote: (1) ComM_Vc_ ^L455^ and (2) ComM_Vc_ ^R497^.

**Fig S7.**
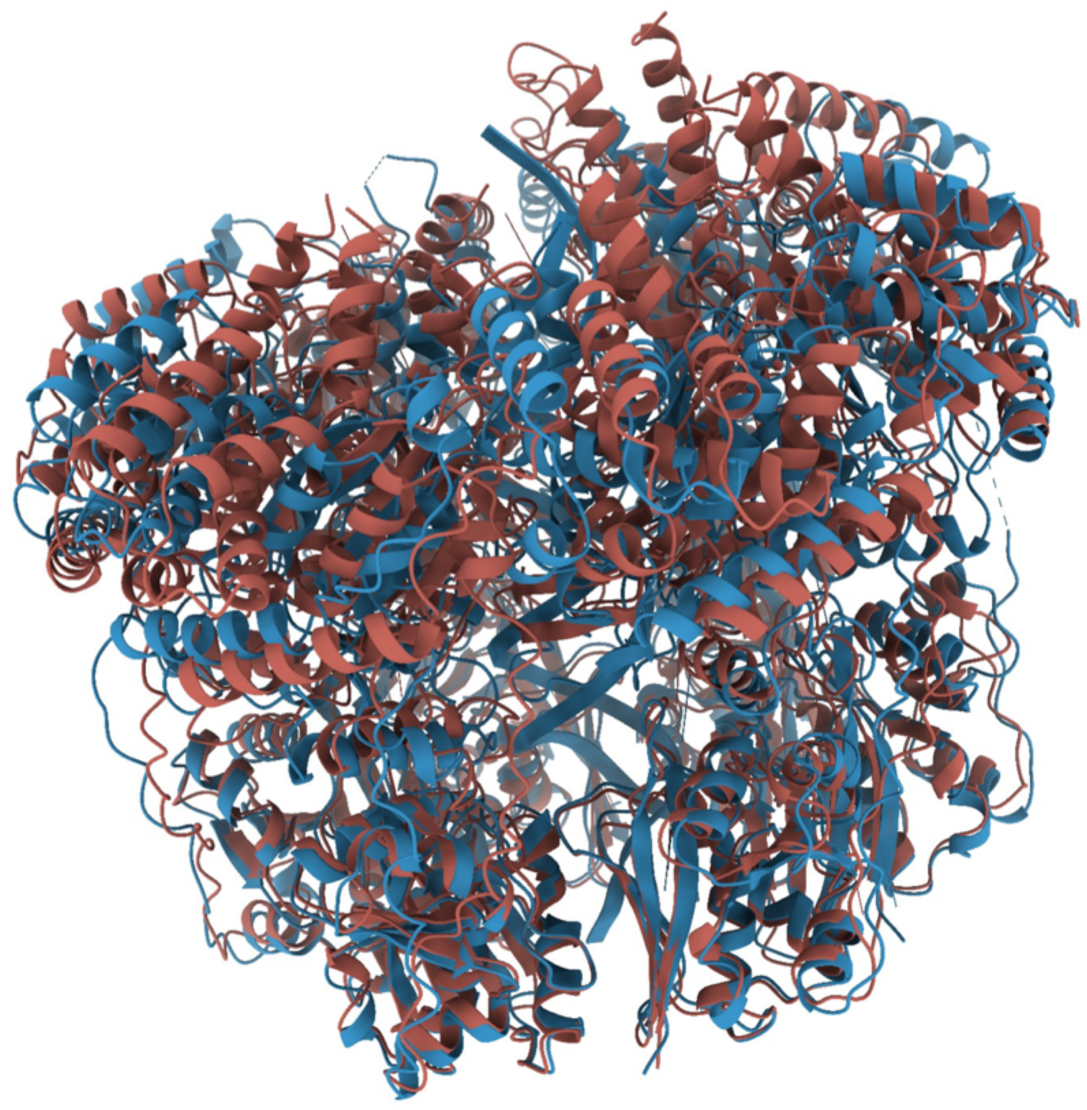
The AF3 ComM model closely aligns to the experimentally derived structure. The AF3 model of the ComM_Vc_ hexamer (brick red) was aligned with the experimentally derived ComM_Lp_ hexamer (blue; PDB: 8RXD). DNA (in blue) is from the experimental structure (PDB: 8RXD). RMSDs were calculated using ChimeraX: RMSD is 2.6 Å for the best pair of ComM monomers (over 375 residues, global RMSD over all 484 residues is 4.3 Å); RMSD is 2.6 Å for the ComM hexamer (over 1394 residues, global RMSD over all 2886 residues is 7.9 Å).

**Fig S8.**
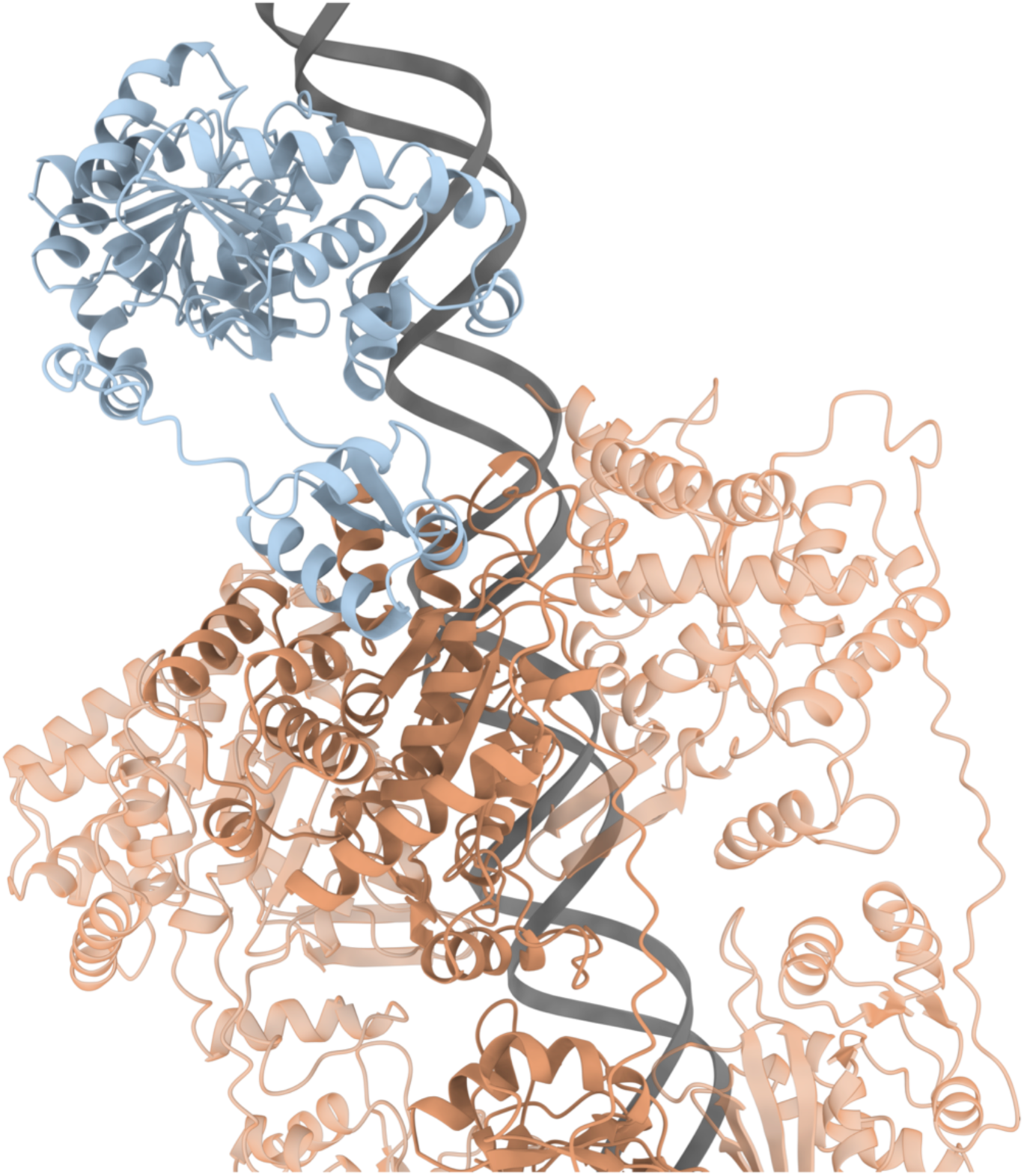
Structural modeling suggests that ComM uses distinct interfaces to hexamerize, interact with DprA, and bind DNA. An AF3 model was generated with 6 x ComM_Vc_ + DprA_Vc_ + double-stranded DNA. Displayed are DprA_Vc_ (blue), 3 ComM monomers for clarity (orange), and the double-stranded DNA.

**Table S1.**
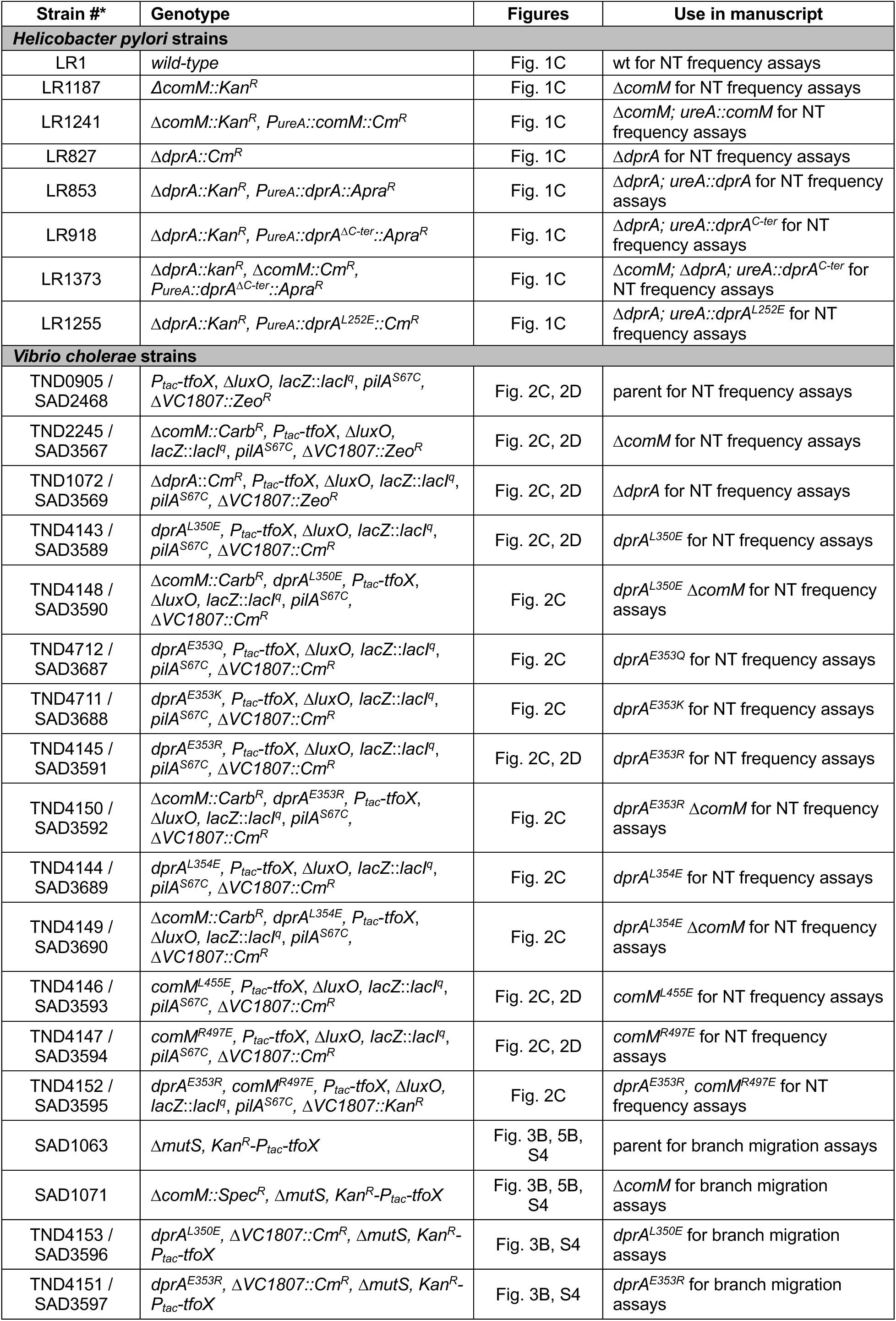
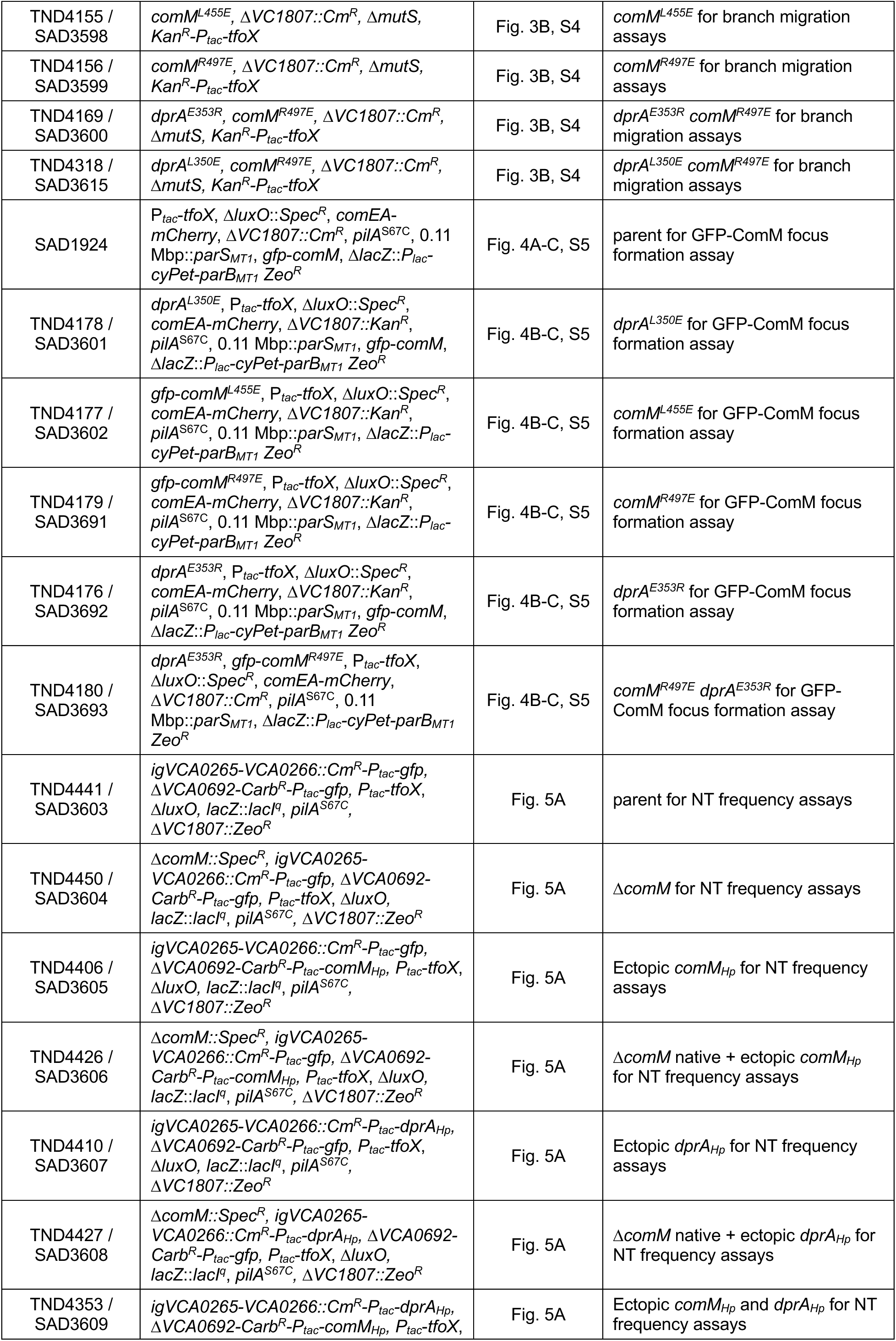
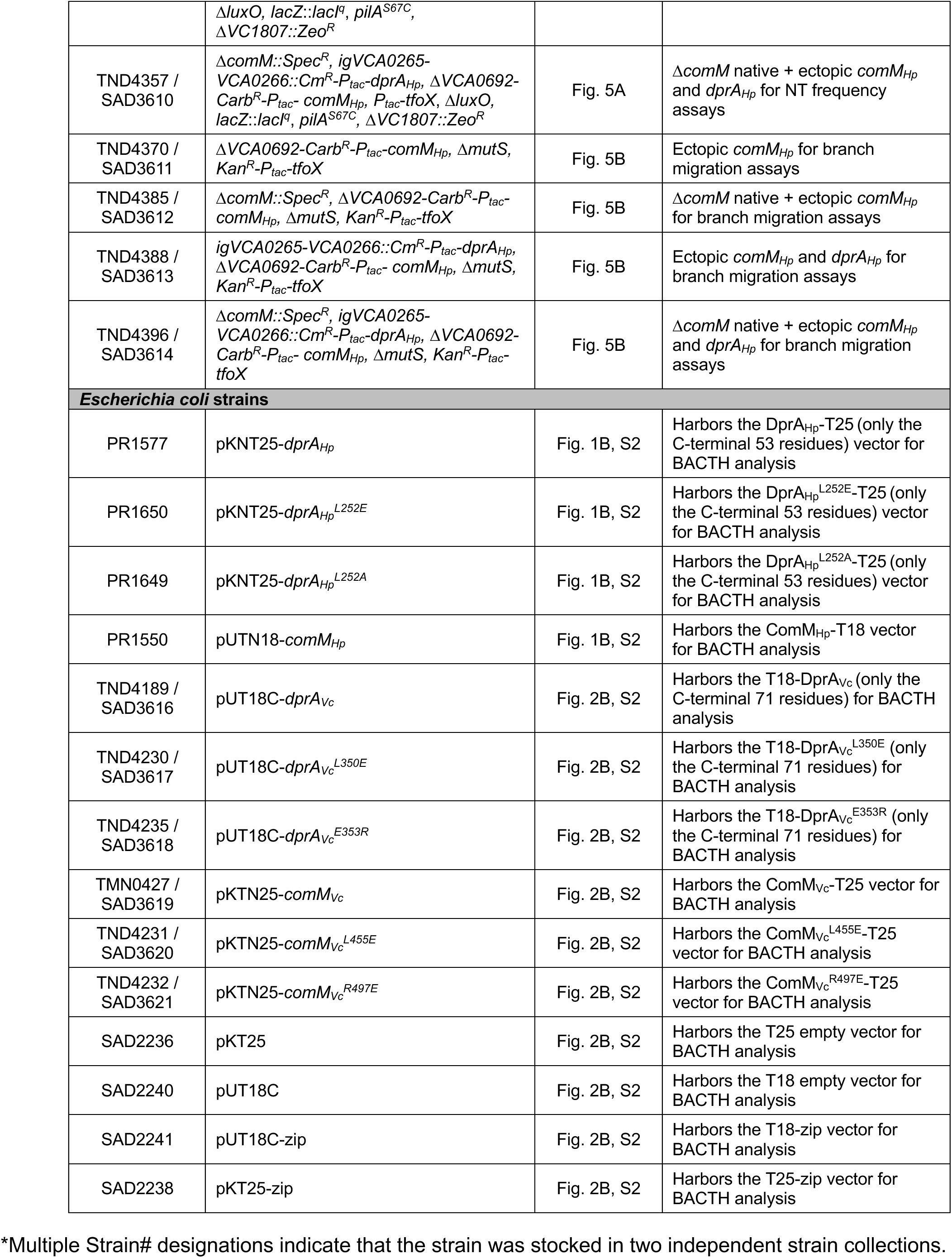
Strains used in this study.

**Table S2.**
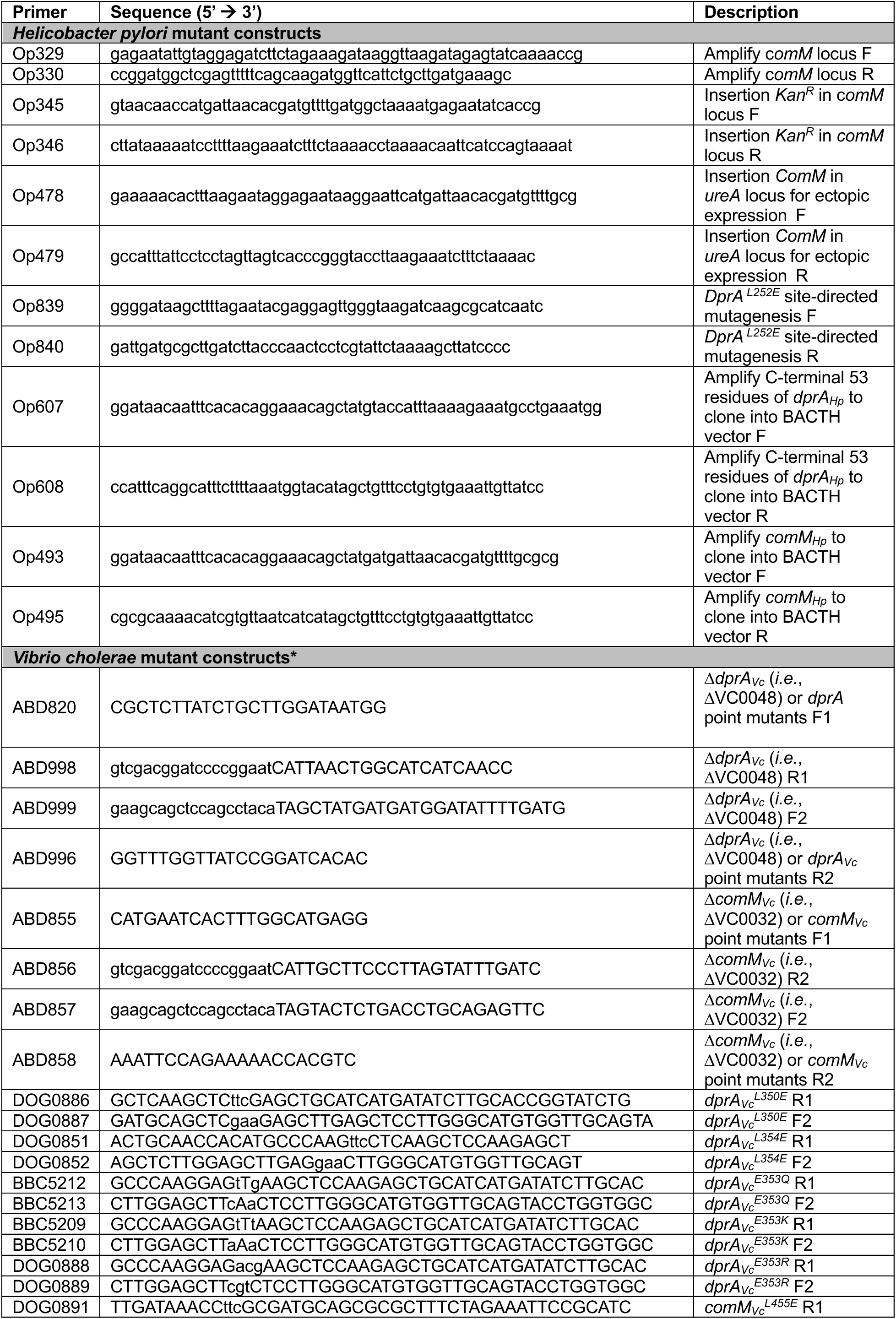
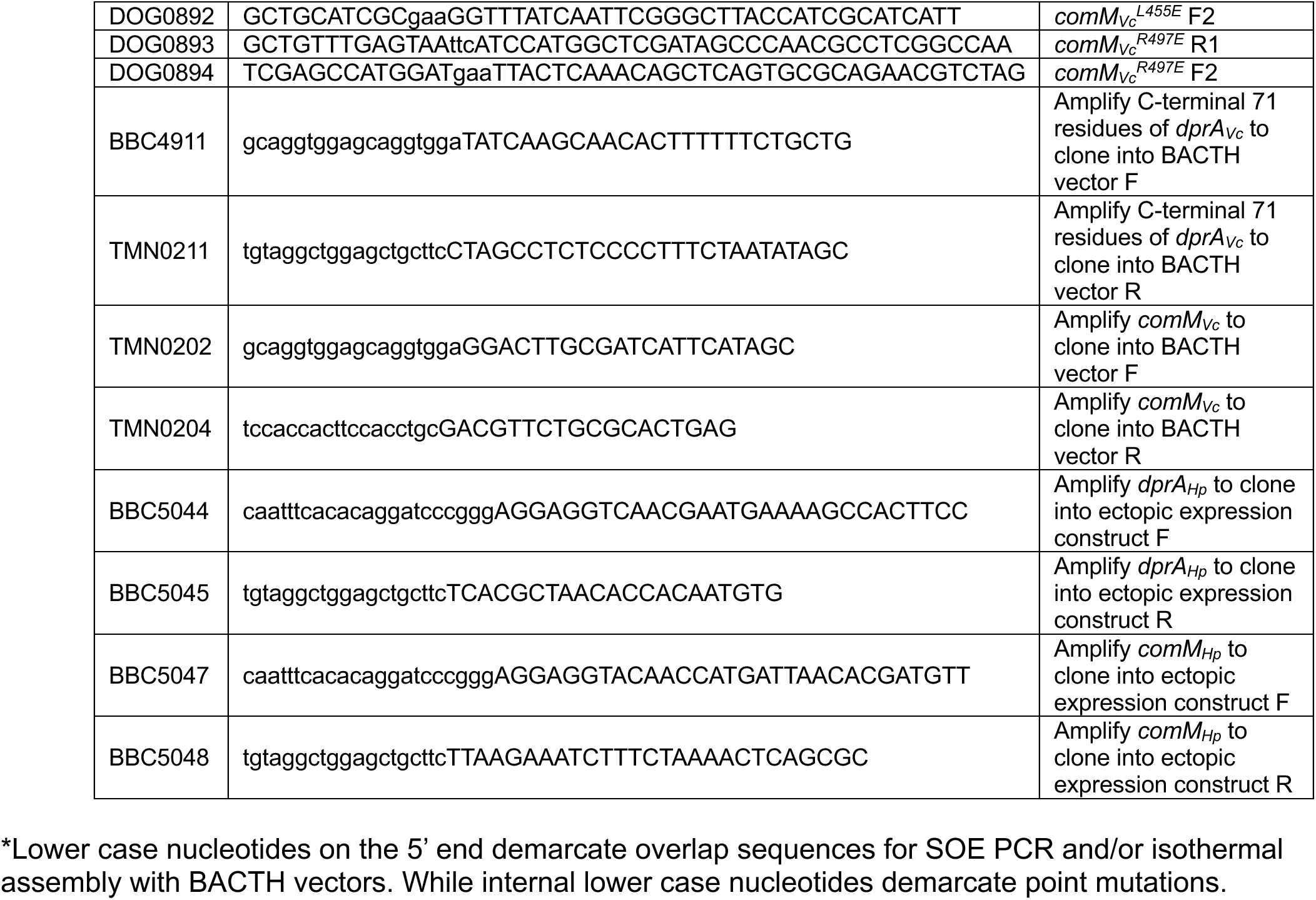
Primers used in this study.

## REFERENCES

1 Dubnau, D. & Blokesch, M. Mechanisms of DNA Uptake by Naturally Competent Bacteria. Annu Rev Genet 53, 217–237 (2019).

2 Johnston, C., Martin, B., Fichant, G., Polard, P. & Claverys, J. P. Bacterial transformation: distribution, shared mechanisms and divergent control. Nat Rev Microbiol 12, 181–196 (2014).

3 Fox, M. S. & Allen, M. K. On the Mechanism of Deoxyribonucleate Integration in Pneumococcal Transformation. Proc Natl Acad Sci U S A 52, 412–419 (1964).

4 Dubnau, D. & Davidoff-Abelson, R. Fate of transforming DNA following uptake by competent Bacillus subtilis. I. Formation and properties of the donor-recipient complex. J Mol Biol 56, 209–221 (1971).

5 Mejean, V. & Claverys, J. P. Use of a cloned DNA fragment to analyze the fate of donor DNA in transformation of Streptococcus pneumoniae. J Bacteriol 158, 1175–1178 (1984).

6 Mortier-Barriere, I. et al. A key presynaptic role in transformation for a widespread bacterial protein: DprA conveys incoming ssDNA to RecA. Cell 130, 824–836 (2007).

7 Lisboa, J. et al. Molecular determinants of the DprA-RecA interaction for nucleation on ssDNA. Nucleic Acids Res 42, 7395–7408 (2014).

8 Quevillon-Cheruel, S. et al. Structure-function analysis of pneumococcal DprA protein reveals that dimerization is crucial for loading RecA recombinase onto DNA during transformation. Proc Natl Acad Sci U S A 109, E2466–2475 (2012).

9 Yadav, T. et al. Bacillus subtilis DprA recruits RecA onto single-stranded DNA and mediates annealing of complementary strands coated by SsbB and SsbA. J Biol Chem 288, 22437–22450 (2013).

10 Bakhlanova, I. et al. Mechanisms of RecA filament nucleation on ssDNA by the DprA protein. bioRxiv, 2024.2005.2007.592916 (2024).

11 Nero, T. M. et al. ComM is a hexameric helicase that promotes branch migration during natural transformation in diverse Gram-negative species. Nucleic Acids Res 46, 6099–6111 (2018).

12 Hardy, L. et al. YraN is a helicase-associated nuclease fostering extended recombination events by natural transformation. bioRxiv, 2024.2002.2006.579203 (2024).

13 Lisboa, J. et al. The C-terminal domain of HpDprA is a DNA-binding winged helix domain that does not bind double-stranded DNA. FEBS J 286, 1941–1958 (2019).

14 Dwivedi, G. R. et al. Insights into the Functional Roles of N-Terminal and C-Terminal Domains of Helicobacter pylori DprA. PLoS One 10, e0131116 (2015).

15 Abramson, J. et al. Accurate structure prediction of biomolecular interactions with AlphaFold 3. Nature 630, 493–500 (2024).

16 Karimova, G., Pidoux, J., Ullmann, A. & Ladant, D. A bacterial two-hybrid system based on a reconstituted signal transduction pathway. Proc Natl Acad Sci U S A 95, 5752–5756 (1998).

17 Dalia, A. B. & Dalia, T. N. Spatiotemporal Analysis of DNA Integration during Natural Transformation Reveals a Mode of Nongenetic Inheritance in Bacteria. Cell 179, 1499–1511 e1410 (2019).

18 Gwinn, M. L., Ramanathan, R., Smith, H. O. & Tomb, J. F. A new transformation-deficient mutant of Haemophilus influenzae Rd with normal DNA uptake. J Bacteriol 180, 746–748 (1998).

19 Rosa, L. T., Vernhes, E., Soulet, A. L., Polard, P. & Fronzes, R. Structural insights into the mechanism of DNA branch migration during homologous recombination in bacteria. EMBO J (2024).

20 Mirouze, N. et al. Direct involvement of DprA, the transformation-dedicated RecA loader, in the shut-off of pneumococcal competence. Proc Natl Acad Sci U S A 110, E1035–1044 (2013).

21 Humphreys, I. R. et al. Protein interactions in human pathogens revealed through deep learning. Nat Microbiol 9, 2642–2652 (2024).

22 Gao, M., Nakajima An, D., Parks, J. M. & Skolnick, J. AF2Complex predicts direct physical interactions in multimeric proteins with deep learning. Nat Commun 13, 1744 (2022).

23 Miller, V. L., DiRita, V. J. & Mekalanos, J. J. Identification of toxS, a regulatory gene whose product enhances toxR-mediated activation of the cholera toxin promoter. J Bacteriol 171, 1288–1293 (1989).

24 Tomb, J. F. et al. The complete genome sequence of the gastric pathogen Helicobacter pylori. Nature 388, 539–547 (1997).

25 Dalia, A. B. Natural Cotransformation and Multiplex Genome Editing by Natural Transformation (MuGENT) of Vibrio cholerae. Methods Mol Biol 1839, 53–64 (2018).

26 Dalia, A. B., McDonough, E. & Camilli, A. Multiplex genome editing by natural transformation. Proc Natl Acad Sci U S A 111, 8937–8942 (2014).

27 Dalia, T. N., Chlebek, J. L. & Dalia, A. B. A modular chromosomally integrated toolkit for ectopic gene expression in Vibrio cholerae. Sci Rep 10, 15398 (2020).

28 Dalia, A. B. & Dalia, T. N. Horizontal gene transfer by natural transformation promotes both genetic and epigenetic inheritance of traits. bioRxiv, 10.1101/596379 (2019).

29 Li, M. Z. & Elledge, S. J. SLIC: a method for sequence- and ligation-independent cloning. Methods Mol Biol 852, 51–59 (2012).

30 Altschul, S. F., Gish, W., Miller, W., Myers, E. W. & Lipman, D. J. Basic local alignment search tool. J Mol Biol 215, 403–410 (1990).

31 Jumper, J. et al. Highly accurate protein structure prediction with AlphaFold. Nature 596, 583–589 (2021).

32 Mirdita, M. et al. ColabFold: making protein folding accessible to all. Nat Methods 19, 679–682 (2022).

33 Steinegger, M. & Soding, J. MMseqs2 enables sensitive protein sequence searching for the analysis of massive data sets. Nat Biotechnol 35, 1026–1028 (2017).

34 Paysan-Lafosse, T. et al. InterPro in 2022. Nucleic Acids Res 51, D418–D427 (2023).

35 Steinegger, M. et al. HH-suite3 for fast remote homology detection and deep protein annotation. BMC Bioinformatics 20, 473 (2019).

36 Ellison, C. K. et al. Retraction of DNA-bound type IV competence pili initiates DNA uptake during natural transformation in Vibrio cholerae. Nat Microbiol 3, 773–780 (2018).

37 Dalia, T. N. & Dalia, A. B. SbcB facilitates natural transformation in Vibrio cholerae in an exonuclease-independent manner. J Bacteriol, e0041924 (2024).

38 Katoh, K. & Standley, D. M. MAFFT multiple sequence alignment software version 7: improvements in performance and usability. Molecular biology and evolution 30, 772–780 (2013).

39 Waterhouse, A. M., Procter, J. B., Martin, D. M., Clamp, M. & Barton, G. J. Jalview Version 2--a multiple sequence alignment editor and analysis workbench. Bioinformatics 25, 1189–1191 (2009).

40 Meng, E. C. et al. UCSF ChimeraX: Tools for structure building and analysis. Protein Sci 32, e4792 (2023).

